# Molecular Mimicry as a Mechanism of Viral Immune Evasion and Autoimmunity

**DOI:** 10.1101/2024.03.08.583134

**Authors:** Cole Maguire, Chumeng Wang, Akshara Ramasamy, Cara Fonken, Brinkley Morse, Nathan Lopez, Dennis Wylie, Esther Melamed

**Affiliations:** The University of Texas at Austin, Department of Neurology; The University of Texas at Austin, Center for Biomedical Research Support

## Abstract

Mimicry of host protein structures (“molecular mimicry”) is a common mechanism employed by viruses to evade the host’s immune system. To date, studies have primarily evaluated molecular mimicry in the context of full protein structural mimics. However, recent work has demonstrated that short linear amino acid (AA) molecular mimics can elicit cross-reactive antibodies and T-cells from the host, which may contribute to development and progression of autoimmunity. Despite this, the prevalence of molecular mimics throughout the human virome has not been fully explored. In this study, we evaluate 134 human infecting viruses and find significant usage of linear mimicry across the virome, particularly those in the herpesviridae and poxviridae families. Furthermore, we identify that proteins involved in cellular replication and inflammation, those expressed from autosomes, the X chromosome, and in thymic cells are over-enriched in viral mimicry. Finally, we demonstrate that short linear mimicry from Epstein-Barr virus (EBV) is significantly higher in auto-antibodies found in multiple sclerosis patients to a greater degree than previously appreciated. Our results demonstrate that human-infecting viruses frequently leverage mimicry in the course of their infection, point to substantial evolutionary pressure for mimicry, and highlight mimicry’s important role in human autoimmunity. Clinically, our findings could translate to development of novel therapeutic strategies that target viral infections linked to autoimmunity, with the goal of eliminating disease-associated latent viruses and preventing their reactivation.

## Introduction

Over the centuries, viruses have co-evolved with their hosts, particularly to gain traits that increase their virulence such as faster replication times, longer infectious periods, and strategies to evade the host immune system^1^. One notable trait is molecular mimicry, in which pathogens “mimic” protein structures of their host^2^. Due to mechanisms generally preventing formation of auto-reactive immune responses^3, 4^, mimicked proteins are thought to be protective to pathogens by limiting the number of pathogenic epitopes able to be targeted by the host, thereby impeding the host’s immune response^2^. However, despite the biological safeguards against autoimmunity, mimicry can elicit T-cell and antibody responses that are cross-reactive to both the mimicking and mimicked proteins, with this cross-reactivity thought to possibly underlie autoimmune pathologies^5–8^.

Previous work has been done to quantify mimicry across a wide range of viral proteomes with an emphasis on determining mimicry at the tertiary and quaternary protein structure level^9^. However, adaptive immune responses generally target small regions of proteins (called epitopes), with T-cells primarily responding to linear protein epitopes of 8-12 or 18-24 amino acids (AAs) long^10–13^, and up to 50% of antibodies binding linear epitopes of 4-12 AAs^14, 15^. Thus, mimicry at the level of primary protein structure (i.e. linear sequence) needs to be evaluated to better understand how viral mimicry may be perceived by the adaptive immune system. While prior studies have evaluated linear AA mimicry in limited cohorts of pathogens^16–18^ or searched specifically for top mimics that may contribute to auto-immunity^19^, it remains unclear how the abundance of linear mimicry differs across a wider range of the human virome.

Interestingly, the “molecular mimicry trade-off hypothesis” posits that mimicry may not always confer a net advantage to the virus^20^. Thus, to adopt mimicry, the advantages gained by the virus must outweigh any associated drawbacks, such as extended replication times due to longer protein sequences or compromised protein functionality due to mutations introduced to achieve mimicry^20^. However, short linear mimicry at the size of an immune epitope may be able to offer substantial evasion of the adaptive immune system while minimizing detrimental effects to either viral protein function or length. As a result, short linear mimicry may possibly offer one optimization to the molecular mimicry trade-off and be advantageous to viruses.

In this study, we conducted one of the largest molecular mimicry screens to date with a total of 134 human-infecting viral proteomes screened for 8mer, 12mer, and 18mer AA fragments with up to 3 mismatches to matching human k-mers. We identify the viral families and strains that have the highest density of mimicry and validate the significance of this mimicry using three permutations tests. Furthermore, we dissect the biological patterns of mimicry and report associated genetic pathways, chromosome location, and cell types. Finally, we evaluate the prevalence of mimicry in the auto-antibodies identified in patients with multiple sclerosis (MS) to demonstrate short linear mimicry might have consequences in autoimmune disorders.

## Methods

### Proteome Retrieval

The full list of human infecting viruses was retrieved from ViralZone^21^ (Table S1) with phylogeny obtained from NCBI Taxonomy^22^. Chronic infecting pathogens were defined as pathogens whose infection on average persists past 21 days (Table S1). All pathogen proteomes used were retrieved from UniProt (Table S1) and compared to the reference human proteome (UP000005640) using suffix array kernel sorting to identify matching 8, 12 or 18 amino acid (AA) fragments (8mer, 12mer, and 18mers) with up to 3 mismatches *in silico* (Figure 1A-C). Only the canonical protein sequences were used for the pathogen proteomes (i.e. no isoforms were used). The 8mer, 12mer, and 18mer length were chosen for their respective association with processing and presentation of peptides on MHC-I (8mer and 12mer) and MHC-II (12mer and 18mer) complexes to CD8+ and CD4+ T-cells respectively^10–13^. In the case of an unknown AA (X) in the human or viral proteome, it was conservatively called a mismatch to all AAs. Proteins that have been deleted from UniProt as of September 12^th^, 2023 were removed from the previously downloaded data from May 2022. Additionally, duplicated proteins in either the viral or human proteomes were removed. When calculating the percent of 8mer, 12mer, or 18mers that had 3 or less mismatches (Figure 1), a mimic was not counted if both the human and pathogen proteins had the same enzyme commission number or were reported as having the same chemical reaction in Uniprot due to concern of homology between similarly functioning proteins (this filter was only used for Figure 1 analyses). Summarized data and statistical results comparing the rate of 8mer, 12mer, and 18mers are reported in Table S2.

**Figure 1.**
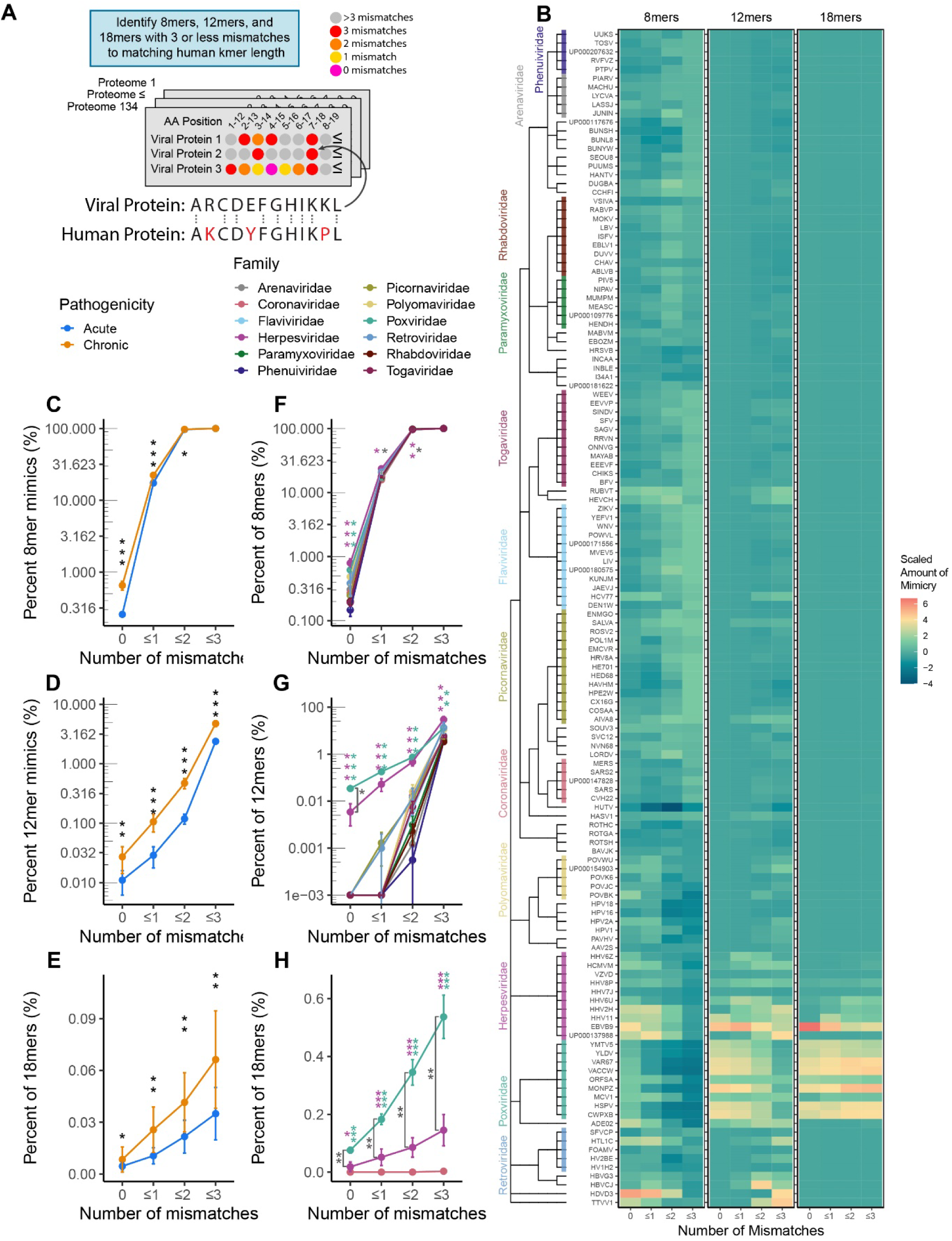
Molecular mimicry across human infecting viruses. **A)** Experimental Schema and example of 12mer AA proteomic alignment. In total, 134 human-infecting viral proteomes were screened to identify 8mer, 12mer, and 18mer AA k-mers with 3 or less mismatches to a matching human k-mer. **B)** Heatmap of scaled percentages of mimicry for 8mers, 12mers, and 18mers with 3 or less mismatches, with viruses aligned by taxonomy. Percents of k-mers, were scaled separetly for each mismatch and k-mer length combination. Percentage of viral **C)** 8mers, **D)** 12mers, and **E)** 18mers with 0, ≤1, ≤2, and ≤3 mismatches between acute and chronically infecting viruses. Percentage of viral **F)** 8mers, **G)** 12mers, and **H)** 18mers with 0, ≤1, ≤2, and ≤3 mismatches between different viral families. Viral familes only shown in F-H if containing at least 5 species. (Black stars indicate comparisons between chronic and acute viruses, magenta and green stars indicate comparisons of the Herpesviruses and Poxviruses against all other viruses, dark gray stars indicate comparison between Herpesviruses and Poxviruses. Kruskal-Wallis test was used for multigroup comparison, and Wilcoxon rank-sum test for pairwise comparison. Error bars denote mean ± standard error of the mean. All p values adjusted for multiple hypothesis testing using Benjamini-Hochberg corrections (* p.adj ≤ 0.05, ** p.adj ≤ 0.01, *** p.adj ≤ 0.001)).

### Permutation Testing of 12mers

To evaluate whether the rate of 12mers was above the expected random chance three permutation strategies were utilized. The first strategy randomly shuffled all proteins in each viral proteome 30 times and fold change (FC) was calculated between the actual proteome and the average of the 30 shuffles. The second strategy reversed all proteins in each viral proteome, for which we calculated the FC over the actual proteome. The third strategy was a quasi-random shuffle with AA shuffled by class defined in Figure 2A, in which all proteins in each viral proteome were shuffled 30 times and FC was calculated of the actual proteome to the average of the 30 shuffles. Statistical testing results for Figure 2 are provided in Table S3.

**Figure 2.**
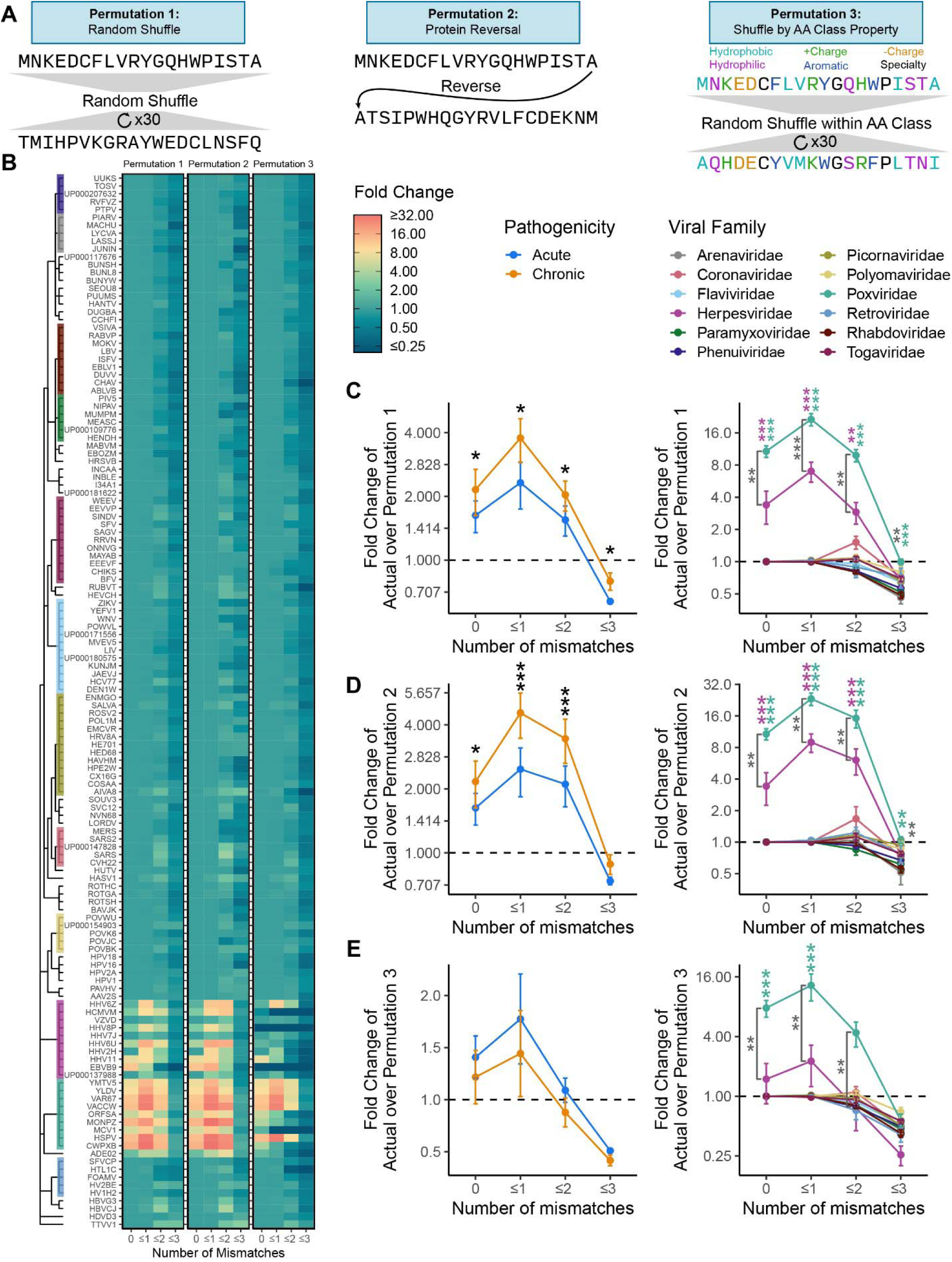
Permutation confirms significant mimicry in poxviruses and herpesviruses. **A)** Example schema of the three permutation strategies (random, protein reversal, AA class shuffle). **B)** A heatmap of the fold change of actual mimicry over mimicry in permutations 1-3, aligned by viral taxonomy. Results of the **C)** random permutation (permutation 1), **D)** protein reversal permutation (permutation 2), and **E)** AA class shuffle permutation (permutation 3) for chronic vs acute viruses and by viral family. Viral familes only shown in C-E if containing at least 5 species. (Black stars indicate comparisons between chronic and acute viruses, magenta and green stars indicate comparisons of the Herpesviruses and Poxviruses against all other viruses, dark gray stars indicate comparison between Herpesviruses and Poxviruses. Kruskal Wallis test was used for multigroup comparison, and Wilcoxon rank-sum test for pairwise comparison. Error bars denote mean ± standard error of the mean. All p values adjusted for multiple hypothesis testing using Benjamini-Hochberg corrections (* p.adj ≤ 0.05, ** p.adj ≤ 0.01, *** p.adj ≤ 0.001)).

### Enrichment Analysis of 12mers

The hypergeometric test was used to evaluate significantly enriched KEGG (v 109.0) pathways mimicked by individual pathogen proteomes using the R package clusterProfiler^23^. Only pathways significantly mimicked by at least three viruses are shown in Figure 4, with all results shown in Figure S4 and reported in Table S4. Enrichment of cell type and cellular location of proteins used data from the human protein atlas (v 22.0), with results reported in Figure S5^24^. Enrichment of mimics to proteins from specific chromosomes and from genes expressed by human mTEC cells in the thymus was evaluated with the Fisher’s exact test, using genes reported in Gabrielsen et al. 2019 with results reported in Table S5^25^.

### Analysis of Phip-seq Autoantibodies

Data was retrieved from Zamecnik et al. 2023^26^. Reads per 100,000 (rpK) were used to calculate FC for each peptides of MS patients based on the control mean and distribution separately for the pre-diagnosis and post-diagnosis timepoint. Peptides with FC > 10 were called as hits.

Suffix array kernel sorting was used against the Phip-seq peptide library to identify 12mers with 3 or less mismatches and 8mers with 3 or less mismatches (following the same protocol as used for the proteome analysis except using the specific human proteome used in the production of the Phip-seq library). The rate of 12mer mimics in “hit” peptides was quantified and compared to non-hit peptides and correlated with prevalence in all samples.

### Statistical Analysis

Computational analyses were performed using the Biomedical Research Computing Facility at UT Austin, Center for Biomedical Research Support RRID#: SCR\_021979 and Texas Advance Computing Center (TACC) The University of Texas at Austin. K-mer alignment was conducted using Julia 1.6.0^27^. All other analyses were conducted using R 4.3.1^28^. The non-parametric tests of Kruskal-Wallis for multiple group comparisons and Wilcoxon rank-sum test for pairwise group comparisons were used for all testing unless otherwise stated. All p-values were adjusted using Benjamini-Hochberg multiple test correction and are referred to as “p.adj” or “adjusted p.values”.

### Data and code availability

Publicly available datasets used in this study are reported in Table 1. All intermediate data and code used in this study are available at https://github.com/melamedlab/MolecularMimicry.

**Table 1.** Key datasets and resources used in this study.

| Dataset/Key Resource | Citation | Version | Notes |
| --- | --- | --- | --- |
| Viral Zone:<br>List of Human infecting<br>viruses | Masson et al.<sup>21</sup> |  |  |
| NCBI Taxonomy:<br>Phylogenetic Information<br>of Pathogens | Schoch et al.<sup>22</sup> |  |  |
| KEGG:<br>Biological Pathway Gene<br>Sets | Kanehisa and Goto<sup>23</sup> | Release 109.0<br>(2024/02) |  |
| Human Protein Atlas:<br>Cell Type, Tissue, and<br>Organ System Protein<br>Expression | Uhlén et al.<sup>24</sup> | Version 22.0<br>[2022-12-07] | Tissue expression was<br>further grouped into organ<br>system with this mapping<br>provided with the source<br>code. |
| Uniprot:<br>Pathogen and Human<br>Proteomes | UniProt Consortium<sup>29</sup> |  | See Table S1 for full list of<br>proteomes used. |
| List of EBV Latent Proteins | Kang et al.<sup>30</sup> |  | Table 1 of Kang et al.<sup>30</sup> |
| Human Thymus Gene<br>Expression Data set | Gabrielsen et al.<sup>25</sup> |  | Table S3 from Gabrielsen<br>et al.<sup>25</sup> |
| Multiple Sclerosis Phip-seq<br>Dataset | Zamecnik et al.<sup>26</sup> |  |  |

## Results

### Elevated Mimicry in Poxviridae and Herpesviridae

To systematically investigate the rate of molecular mimicry in human infecting viruses, we queried 134 viral proteomes for the matching percentage of amino acid (AA) 8mers, 12mers, and 18mers between human and viral proteins with up to three mismatches (Figure 1A). The evaluated AA length (k-mer) was motivated by the known length of T-cell recognized epitopes which range on average from 8-12 AA for CD8+ T-cells and 18-24 AA for CD4+ T-cells^10–13^. We found that the phylogenetic relationship between pathogens and the relative amount of mimicry was similar for 8mers, 12mers, and 18mers between closely related viral species (Figure 1B).

In comparing acute and chronically infecting viruses (infection less than or greater than 21 days respectively), chronically infecting viruses had significantly elevated rates of mimicry for 0, ≤1, and ≤2 mismatches for all evaluated k-mers (8, 12, and 18 AAs) and for ≤3 mismatches in 12 and 18mers (Figure 1C-E). On further evaluation by viral family, while both Herpesviridae and Poxviridae had significantly higher levels of 8mer mimicry at 0 mismatches, only Herpesviridae demonstrated significance for ≤1 and ≤2 mismatches (Figure 1F). In turn, for 12mer and 18mers, Herpesviridae and Poxviridae were both significant for 0, ≤1, ≤2, and ≤3 mismatches compared to all other viral families (Figure 1G-H). Notably, Poxviridae had significantly more 18mers with 0, ≤1, ≤2, and ≤3 mismatches compared to Herpesviridae (Figure 1H), suggesting that Poxviridae may be more adept at mimicking longer AA sequences. In line with these findings, we also observed that classifying viruses by genome structure (Baltimore classification, e.g. dsDNA, ssDNA, +/-ssRNA) revealed elevated rates of mimicry in dsDNA viruses (driven largely by Herpesviruses and Poxviruses, Figure S1).

Interestingly, the observed elevation in mimicry across diverse viruses was not limited to specific viral proteins (e.g. known virokines) but rather was more broadly distributed throughout the proteomes (Figure S2A-B). Furthermore, relative amino acid usage did not greatly differ between viruses and was unlikely to explain the significant differences in mimicry (Figure S2C).

### Permutation Testing reveals Significant Rates of Mimicry in Herpesviruses and Poxviruses

To identify whether the rate of detected mimicry was above random chance, we conducted three permutation tests using 12mers: i) scrambled viral proteins (permutation 1), ii) reveresed viral proteins (permutation 2), and iii) scrambled viral proteins by amino acid (permutation 3)(Figure 2A-B). We chose to utilize 12mers over 8 and 18mers for two reasons: first, most viral 8mers had a human 8mer match at ≤2 mismatches and second, 18mers were too sparse for detection across most viral families (Figure 1F and 1H). When the actual proteomes were compared to the mean of the randomly scrambled proteomes, chronic viruses and the specific families of Poxvirdae and Herpesvirdae were confirmed to have significant levels of mimicry at 0, ≤1, and ≤2 mismatches (Figure 2C). However, this permutation strategy does not fully capture the fact that proteins form secondary structures that affect the underlying AA sequence (e.g. alpha helices, beta-pleated sheets). To address this gap, we next evaluated viral proteomes with reversed protein sequences, with this strategy better maintaining secondary structures in the permutations. This permutation also confirmed that the rate at which we observed viral 12mers with 0, ≤1, and ≤2 mismatches was significantly above random chance in chronic viruses and specifically in Poxviridae and Herpesviridae (Figure 2D). Finally, as the most stringent permutation, we scrambled AAs by class (e.g. hydrophobic, hydrophilic, postively charged, etc), which conserved overall protein structure of AA properties. We observed a similar trend to the first two permutations, with Herpesviridae significant for 12mers with 0 mismatches and Poxviridae significant for 12mers with 0 and ≤1 mismatches (Figure 2E). The weakened significance compared to the first two permutations likely reflects the overly stringent nature of this permutation as several AAs were intentionally not shuffled (e.g. Cystine, Glycine, and Proline), and AAs in each class ended up shuffled back into their original position at a higher frequency compared to the first two permutations.

### Characteristics of Peptide Mimicking by Viral Family

Having confirmed mimicry was significantly higher in Herpesviridae and Poxviridae, we next evaluated the AA length of observed mimicry. Due to the nature of the rolling window approach used in the alignment, we could identify when several k-mers in a row aligned to the same protein resulting in a k-mer “run” (example shown in Figure 3A). These runs allowed us to identify how long the potential mimicry sequence could be (e.g. three 12mers in a row for a single protein forming a single 14mer, five 12mers in a row forming a single 16mer). Using this approach, several viral families were identified as having longer runs on average including Herpesviridae and Poxviridae (Figure 3B, Figure S3). Notably, there were several very long k-mers (>30 AA) that reflect known full protein mimics such as vIL-10, a viral analog of an anti-inflammatory cytokine^31^.

**Figure 3.**
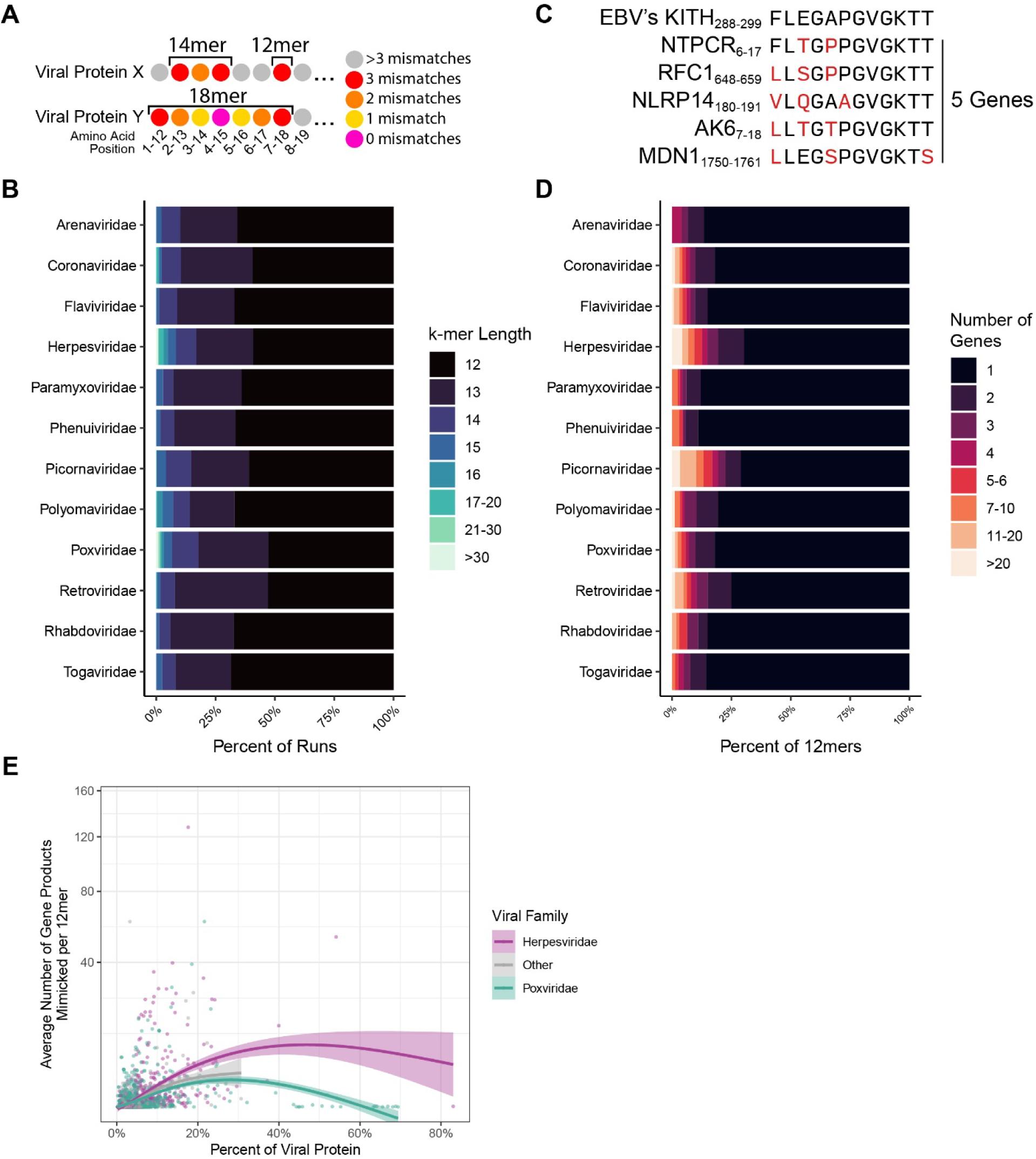
Viral mimicry is diverse in length and differentially targets human protein motifs. **A)** An example of an 18mer and 14mer comprised of 12mers. **B)** Average percent of mimics at varying k-mer lengths. **C)** An example of multi-mapping in which a single viral 12mer aligns to multiple human proteins with the criteria of 3 or less mismatches. **D)** Percent of viral 12mers and their corresponding number of mimicked human genes (multi-mapping). **E)** Average number of host gene products mimicked, plotted against the percent of the viral protein that participates in mimicry (defined as a 12mer with 3 or less mismatches).

Additionally, we also identified some viral families that had more “multi-mimics” where a single viral 12mer had more than one human 12mer with 3 or less mismatches (Figure 3C-D). This result reflects that these 12mers are mimicking a human motif scattered across multiple genes that may perhaps be more advantageous to mimic. Interestingly, when evaluating the mean rate of multi-mimics versus the percentage of viral 12mers participating in mimicry (i.e. percent of 12mers with 3 or less mismatches to a human 12mer) for each viral protein from across all proteomes in the cohort, we observed that multi-mimics are more frequently found in proteins with lower percentages of mimicry. Notably, overall Herpesviridae proteins display a greater degree of multi-mimicking (Figure 3E).

### Viral mimics target key human cellular pathways

Next, we evaluated whether the human mimicked proteins were overenriched in specific biological pathways. Using the KEGG database^23^, we identified several pathways that were significantly enriched across different viral proteomes (Figure 4A), with the pathways primarily relating to cellular replication and inflammation (Figure 4B and S4). Notably, many pathways’ significance was a result of “multi-mimics” that reflect mimicry of central human motifs in the pathways. When enrichment testing was repeated with only viral mimics that mimicked five or greater different gene products, many pathways were still significant (Figure 4A, indicated with an X).

**Figure 4.**
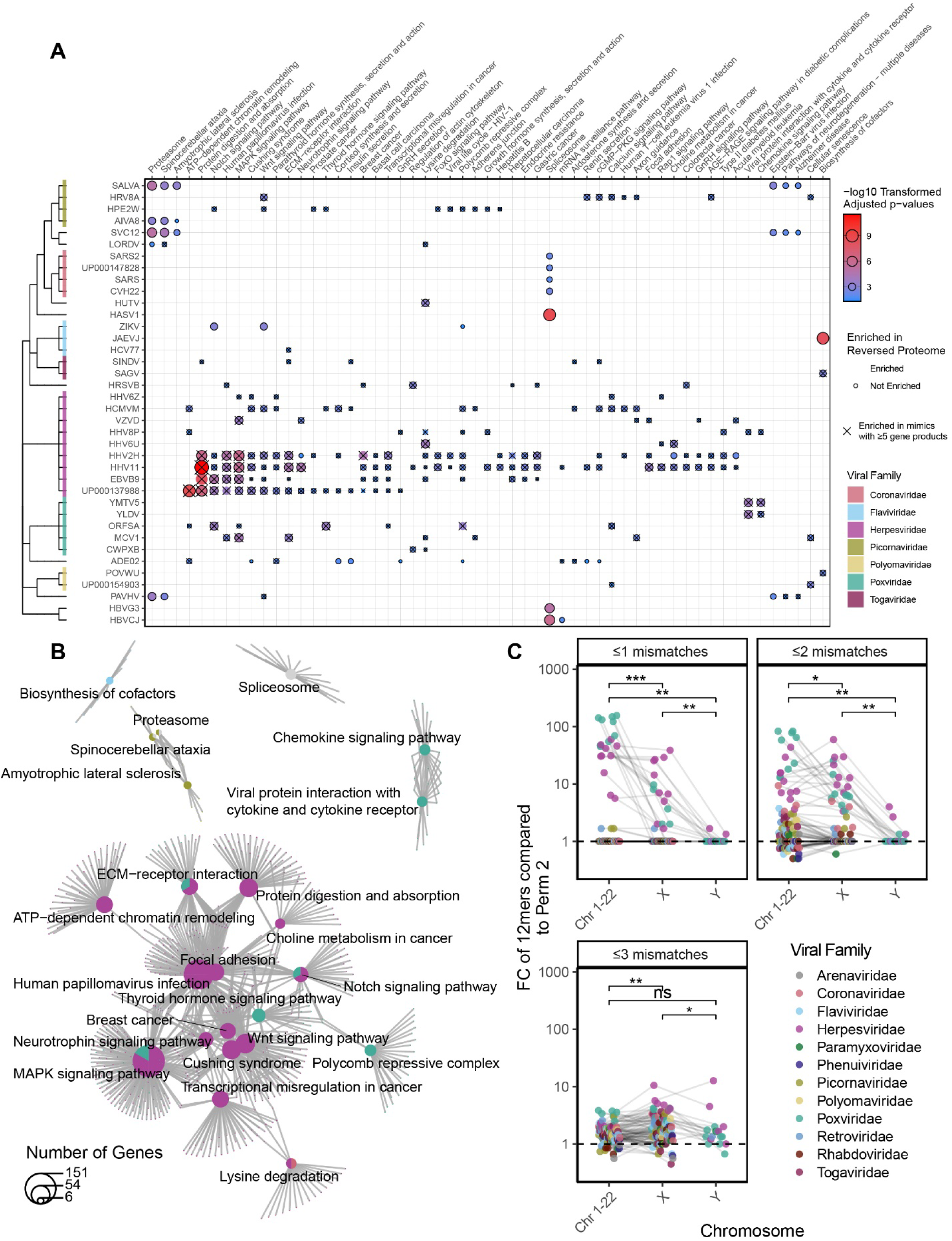
Viral mimicry targets vulnerable biological pathways. **A)** Hypergeometric enrichment testing of KEGG pathways for human proteins that are mimicked by each virus. Significant enrichment is displayed as a dot and is outlined if that biological enrichment was not observed in the reverse proteome permutation (permutation 2). Only pathways significant in at least 3 viruses are shown, with all viruses and significant pathways shown in Figure S4. **B)** Shared overlap between the significant pathway from **(A)** reveal broad roles of inflammation and cellular replication amongst the pathways. Pathways and genes are represented by a pie chart, colored by the proportion of viruses belonging to each family that were significantly enriched for the pathway, with lines connecting genes to their respective pathways. **C)** Fold change of the percent of mimics whose human conterpart is encoded on either an autosome (Chromosomes 1-22), X, or Y chromosome over the rate in the reversed proteome (permutation 2). (Hypergeometric enrichment used for pathway analysis. Wilcoxon summed-rank test used for paired pairwise comparison. All p values adjusted for multiple hypothesis testing using Benjamini-Hochberg corrections (* p.adj ≤ 0.05, ** p.adj ≤ 0.01, *** p.adj ≤ 0.001)).

Leveraging the Human Protein Atlas, we also evaluated the rate at which proteins from specific cell types, tissues, or organ systems were mimicked, though no strong association was observed (Figure S5). Instead, pathogens mimicked from a myriad of proteins non-specific to tissues or cell types, in line with our finding that mimicry was dispersed throughout viral proteomes (Figure S2A-B).

### Viruses’ mimicry targets proteins from autosomes and the X chromosome

To further investigate the potential of evolutionary pressure on viral molecular mimicry, we evaluated chromosome location of the mimicked human proteins. As the Y chromosome is only carried in males, we hypothesized that viruses would display preference for mimicking autosome and X chromosome proteins found in the entire population. Using the reversed proteome as a reference, viruses had significantly greater mimicry to proteins from the autosomes and the X chromsome (Figure 4C). The result was replicated with the randomly shuffled and AA class shuffled permutations (Figure S6).

### Greater viral mimicry in latently expressed proteins of EBV compared to lytic proteins

To analyze whether the timing of protein expression during the viral life cycle (Figure 5A) may explain degree of mimicry, we evaluated one of the top mimicking pathogens, EBV, as expression of its latent and lytic proteins have been well characterized^30^. Utilizing previously reported gene sets^30^, we found that latent stage proteins displayed significantly more mimicry than lytic stage proteins (Figure 5B). This observation was not replicated in HHV8 or CMV, however these viruses still maintained overall high levels of mimicry in both their latent and lytic proteins (Figure S7).

**Figure 5.**
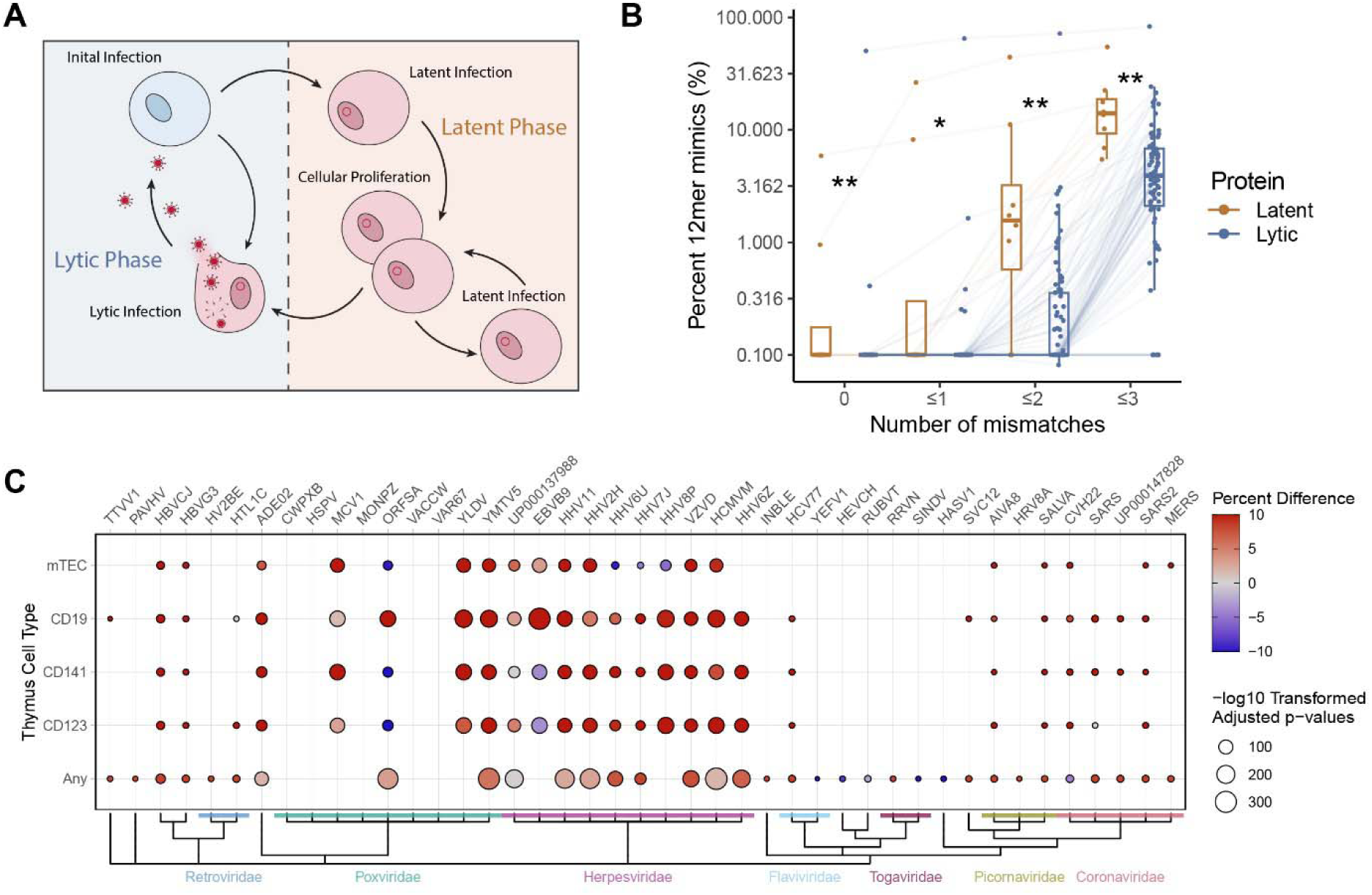
EBV latent proteins have elevated mimicry and viral mimicry targets negatively selected proteins. **A)** Schematic of latent vs lytic stages of viral replication. **B)** Percent of 12mers with 0, ≤1, ≤2, and ≤3 mismatches in latent and lytic EBV proteins. Each point denotes a single EBV protein connected across the mismatch levels with boxplots denoting the interquartile range. **C)** Fisher test results of the proportion of mimics (12mers ≤2 mismatches) to proteins identified previously as expressed in mTEC, CD19+ B-cells, CD141+ dendritic cells, CD123+ dendritic cells, or “any” of these cells in the human thymus (all antigen presenting cells), compared to the proportion of all human proteins expressed by these cell types (as determined by RNA expression in Gabrielsen et al.^25^). Color indicates the percent difference between the percent of mimicked proteins expressed in the thymus and the rate of all human proteins expressed in the thymus.

### Enrichment of Thymic Negatively Selected Proteins

We hypothesized that some viruses may intentionally aquire mimicry to protect their epitopes from being targeted by the adaptive immune system, as thymic negative selection would deplete T cells capable of responding to human mimicked epitopes. To test this hypothesis, we evaluated the proportion of the viral mimics that aligned to genes known to be expressed in human thymus antigen presenting cell populations (mTECs, CD19+ (B cells), CD141+ (dendritic cells), CD123+ (dendritic cells), or “Any” reflecting the list of genes expressed in any of these cell types), as genes not expressed in the thymus during negative selection have more limited mechanisms preventing formation of auto-reactive T-cells (i.e. peripheral tolerance)^32^. Using Gabrielsen et al.’s^25^ reported gene expression of human thymic cells, we observed that the viral mimicking sequences disproportionally mimic proteins found in human thymic cells when compared to the frequency of all human proteins expressed in those cell types (Figure 5C)^25^. Notably, every virus evaluated in the Herpesviridae family maintained a robust relationship with at least one thymic cell type.

### Molecular mimicry explains portions of the multiple sclerosis auto-antibodyome

Finally, we sought to evaluate whether our viral mimics could explain auto-antibodies in a pathological autoimmune disease. Due to the strong epidemiological association between multiple sclerosis (MS) and EBV^33^, we leveraged the recently generated data from Zamecnik et al.^26^ to further evaluate how linear mimicry may play a role in the MS autoimmune pathophysiology. Previously, Zamecnik et al.^26^ screened for auto-antibodies against the human proteome using Phip-seq, a technique used to identify auto-antibodies against 49 AA long linear peptides from the human proteome. The authors found that approximately 8% of MS patients will have an auto-antibody signature that contains a motiff found in EBV’s Envelope Glyocprotein M and BRRF2, with this signature refered to as an immunogenic cluster (IC Cluster).

In line with Zamecnik et al.’s findings, we observed that even non-IC cluster MS auto-antibodies are more likely to contain EBV mimicry (Figure 6A). This association is even stronger for the most prevalent auto-antibodies. In fact, 87% (20/23) of the top MS auto-antibodies found post MS diagnosis outside of the IC-cluster contained an EBV 8mer with ≤2 mismatches compared to 72% in the healthy control auto-antibodies (Figure 6B). Furthermore, we identified that these additional MS auto-antibodies were not fully concomitant with the IC signature, suggesting that EBV mimics between MS patients may differ (Figure S8).

**Figure 6.**
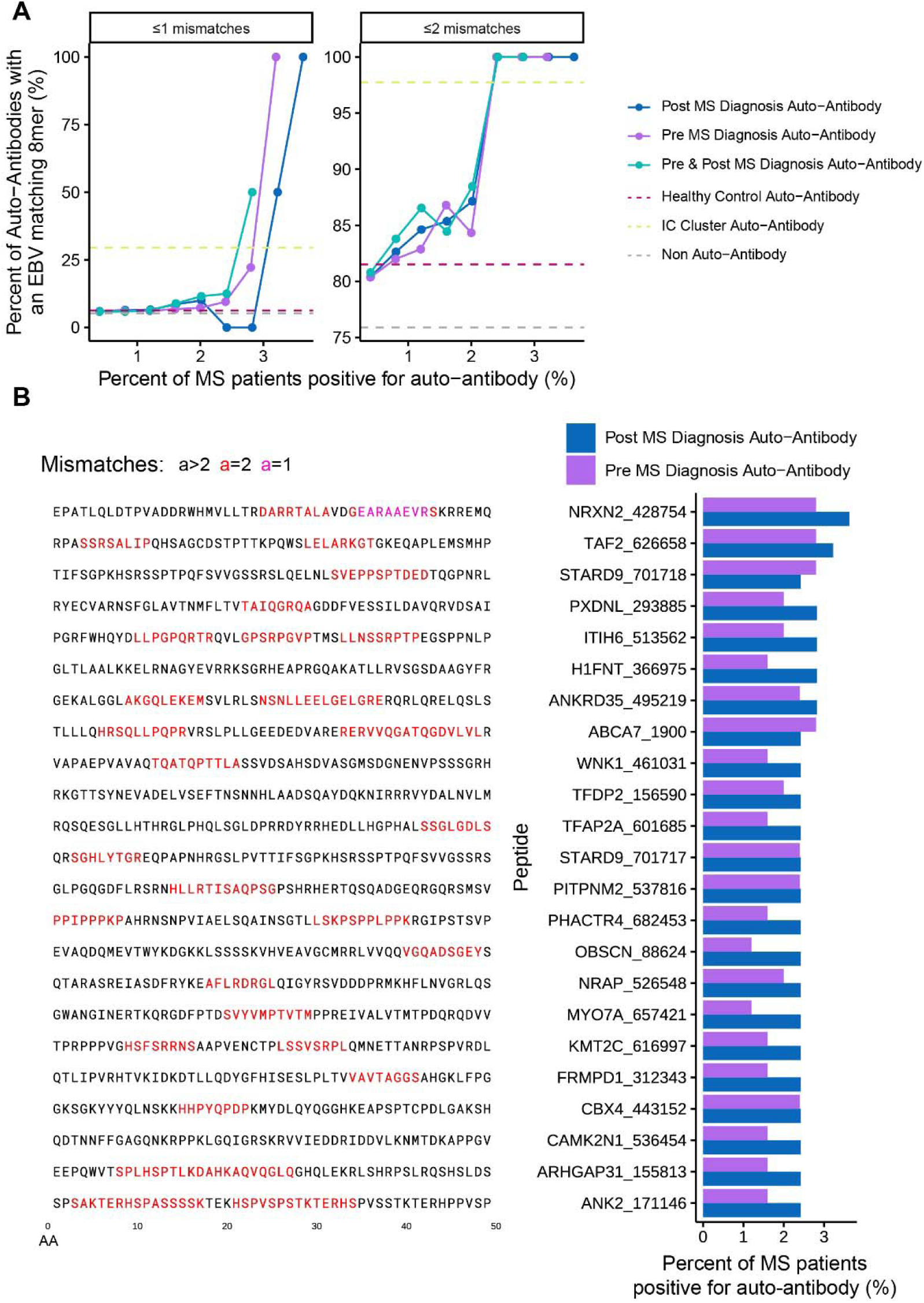
EBV mimics are more frequent in MS auto-antibodies. **A)** Frequency of EBV mimicry (defined as EBV 8mers with ≤1 (left) or ≤2 mismatches (right)) in autoantibody targets in MS patients pre and post diagnosis. Dotted lines show the rate of peptides with an EBV mimic for auto-antibodies found in either in no participannt, in healthy controls, or in the previously identified IC Cluster (Zamecnik et al.^26^). **B)** Left: Peptide sequences of the top non-IC Cluster and most frequent post MS diagnosis auto-antibodies. Right: Percentage of MS patients positive for the top non-IC Cluster auto-antibodies at both the pre and post diagnosis timepoints.

## Discussion

In this study, we evaluated 134 proteomes from all existing human infecting viruses for short linear molecular mimicry to human proteins, comprising one of the largest viral screens to date. These results broaden the current knowledge on the rates of mimicry, as prior studies have primarily focused on limited number of pathogens or mimics predicted to bind to common HLA haplotypes^16–19^. Furthemore, our research reveals underlying cellular and biological patterns associated with mimicry, highlighting the complexity and molecular consequences of host-pathogen interactions. Overall, our results both capture a more holistic picture of viral mimicry and viral evolutionary adaptations relative to prior studies and offer potential avenues for targeted therapeutic strategies in autoimmune diseases.

Notably, we found that viruses within the Herpesviridae (e.g. EBV, CMV, HSV, Varicella Zoster) and Poxviridae (e.g. Monkeypox, Orf virus, Molluscum contagiosum) families have significantly more short linear mimicry relative to all other viruses. Additionally, we demonstrate that the observed elevated rates of mimicry in Herpesviruses and Poxviruses are significantly over-represented compared to permuted forms of their proteomes, suggesting an evolutionary pressure for mimicry. In line with evolutionary pressure, we observe that mimicry to proteins from the autosomes and X chromosome was significantly greater than proteins from the Y chromosome, which would offer less evolutionary benefit to the virus due to only existing in approximately half of the human population. Furthermore, we identified that viruses disproportionately mimicked proteins expressed during negative selection in the thymus (a key biological process in apoptosis of auto-reactive T-cells during their development)^4, 32^, in concordance with our hypothesis that mimicry might improve evasion of the adaptive immune system. In addition, we observe that for one of the most common herpesviruses (EBV), mimicry is significantly higher in latently expressed proteins compared to lytic proteins, further suggesting evasion of the adaptive immune system may be more advantageous during the latent stage of infection. These findings collectively raise an intriguing question about the co-evolution of viruses with the human genome and whether there may be consequences from viral mimicry to the host.

Interestingly, Herpesviridae, which are chronically infecting viruses, have been previously associated with autoimmune diseases (e.g. EBV in MS^33^, CMV in systemic lupus erythematosus ^34^), with molecular mimicry hypothesized as one of the major mechanisms leading to cross-reactive autoimmune responses^5–8^. Similarly, infection with acute respiratory and gastrointestinal viruses has also been observed more frequently in weeks prior to autoimmune disease onset^35–38^; however, the mechanistic link between acute infecting pathogens and autoimmunity remains unknown, unlike with Herpesviridae. Of note, recent work has shown that during infection with COVID-19, ∼41% of patients will have reactivation of EBV, ∼28% HSV1/HSV2, ∼25% CMV, and ∼34% HHV6^39^. Thus, it is possible that herpesviruses, as opportunistic pathogens, may take advantage of other acute infections to exit latency, which may explain why many acute viral infections appear to be epidemiologically associated with autoimmunity.

In comparison, poxviruses have not been associated with autoimmune pathology, despite also displaying significant mimicry. This may be explained by poxviruses mimicking a less diverse number of proteins, focusing instead on functional mimics to modulate host response (e.g. NFKb-I), which often are longer and more accurate mimics. In contrast, herpesviruses display widespread distribution of short and low accuracy mimics (as evident by herpesviruses being significantly higher than poxviruses for 12mers with ≤3 mismatches, but not ≤2 mismatches), which might be sufficient to extort host negative selection to improve evasion of the adaptive immune system. Furthermore, due to establishing a latent infection, the immune system will more frequently encounter and activate against herpetic viruses, possibly increasing the likelihood that a mimicked region is targeted. Conversely, poxviruses are more rarely encountered in the human population and rarely establish latency, thus representing a less frequently encountered infection^40, 41^.

Previous studies on the development of autoreactive T-cells due to mimicry have identified that cross-reactivity may primarily be evoked in non-naive T-cells^42^. It is hypothesized that in the naïve state, the T-cell receptor (TCR) is specific to only the viral peptide, but as the T-cell transitions into a memory state post-infection and develops a lower threshold for activation, the TCR becomes capable of responding to both the viral and the human mimicked peptide. This hypothesis is intriguing as it may explain why cross-reactive T-cells may not be negatively selected during development, and why molecular mimics targeted for autoreactivity often have mismatches in the epitope^43, 44^. Interestingly, this is in line with our observation that herpesviruses have higher levels of short and low accuracy mimicry which may be prime for eliciting autoimmunity.

The high dispersal of mimicry throughout a viruses’ proteome at an overall low frequency is of interest, as it potentially provides insights into mechanisms by which mimicry is acquired and maintained. First, low frequency proteome-wide dispersal of mimicry might better avoid interference with protein structure and function, thus addressing the molecular mimicry trade-off hypothesis^20^. Second, low frequency mimicry also suggests that most of these events are not due to horizontal genetic transfer, except for a few outliers that do reflect horizontal gene acquisition (Figure 3E). Similarly, we did not observe enrichment of mimicry in proteins specific to the tissues and cell types each virus infects, an observation that would otherwise suggest more convincing evidence of horizontal gene transfer. The molecular mimicry trade off hypothesis also identifies that adding mimicry to the viral genome may lengthen replication time, an important trait in overall virulence. Interestingly, we observe that viruses with the most mimicry have some of the largest genomes, possibly indicative of less evolutionary pressure on replication time which might enable them to carry more mimicry. Thus, despite the potential drawbacks of mimicry, its usage on a micro scale appears to be extremely viable.

The observed mimics (12mers) were significantly enriched for key biological pathways, primarily pertaining to cellular replication and immune signaling, suggesting that some of these mimics may be “functional”. Functional mimicry is defined as a pathogenic mimic capable of eliciting complete, partial, or modified effects of the original host protein, with some examples including mimicry of IL10 and NFKB-I, which all further modulate host immune responses^45, 46^. Often, functional mimicry is a byproduct of horizontal genetic transfer and has traditionally been studied in the context of mimicry of complete proteins. However, transcription factor and protein binding sites can comprise small portions of a protein, as small as 4 AAs, which the 12mers in this study would encompass. Thus, mimicry on a smaller scale might also offer some functional effects. Interestingly, the significant enrichment we observe towards cellular replication and immune signaling pathways was often the result of mimicry of key motifs found throughout multiple proteins in the pathway. Thus, some of these mimics might offer functional benefit, particularly in modulating signaling associated with the original mimicked protein.

Finally, we demonstrate the direct translatability of our results to autoimmunity in MS, one of the most common autoimmune diseases^26^. The recent study by Zamecnik et al^26^ detailed an auto-antibody profile in ∼8% of MS cases, believed to stem from EBV-induced molecular mimicry. Here, we extend these findings and demonstrate that even the rare MS auto-antibodies not in this signature, found in only ∼2-4% of MS patients (and absent in controls), also display over enrichment of EBV mimicry, suggesting that EBV mimicry may play a larger role in the auto-antibodyome of MS than previously thought.

Given our data and emerging role of viruses in autoimmune diseases, a better understanding of molecular mimicry may also inform future therapies and interventions in autoimmunity. For example, in MS, clinical interventions for EBV management are currently being evaluated including infusion of EBV specific autologous T-cells and repurposing of anti-virals such as tenofovir/emtricitabine to limit EBV replication^47, 48^. Thus, identifying pathogens as risk factors in specific autoimmune diseases could pave the way for tailored therapeutic strategies that incorporate both the autoimmune disease-specific therapy as well as targeted management of the implicated viral infection(s).

## Limitations

In this manuscript, we specifically focus on linear mimicry, and as a result we may have missed structural 3D mimics, which might be important, particularly to cross-reactive antibody binding. Furthermore, in our evaluation of linear mimicry we calculate the number of mismatches in AA 8mer, 12mer, and 18mers to identify approximate patterns of molecular mimicry. However, mismatches in key positions of the k-mer may differentially affect strength of mimicry, particularly in relation to HLA binding. Yet, the consistent trend of elevated rates of mimicry across all three k-mer lengths and across various levels of mismatches supports the idea that the observed mimicry is robust. Additionally, the evolutionary gain of molecular mimicry was not directly confirmed in this study due to the limited availability of historical data for validation. Thus, future work should leverage ongoing viral monitoring of circulating strains to determine if molecular mimicry is selected for over time. Finally, these results use the consensus viral and human proteomes which do not fully capture genetic diversity in the population. Thus, further examining viral substrains, particularly in relation to the genetics of local populations they infect, may reveal even stronger trends than those observed in this study.

## Conclusion

In conclusion, we performed one of the most comprehensive viral mimicry assessments to date across all known human-infecting viruses, examining short mimicry of peptide lengths commonly recognized by the adaptive immune system. We identified significant levels of mimicry, particularly in the Herpesviridae and Poxviridae families, and report associated biological patterns. Our results provide evolutionary insights into immune evasion and how evolutionary pressures may be shaping viral mimicry. In addition, our results highlight the potential role of viral mimicry in the development of autoimmune diseases like MS, revealing new avenues for therapeutic strategies, such as targeting viral infections linked to autoimmunity, with the goal of mitigating the viral triggers contributing to these diseases.

## Acknowledgements

We appreciate discussions with John Moore, Dr. Victoria Chu, Dr. Lauren Ehrlich, Dr. Laura Fonken, Dr. Dayne Mayfield, and Dr. Hans Hofmann. We thank Advance Microdevices for contributing computational servers used in this study. We are also grateful for the administrative and technical support from Dell Medical School Neurology Department and the UT Austin Biomedical Resource Computational Facility.

## Author Contributions

CM and EM conceived this work. DW and CM conceived the methodology and software. CM, CW, AR, CF, BM, and NL undertook the formal analysis. CW, AR, CF, BM, and NL obtained resources and data curation. CM was responsible for visualization. EM obtained funding. CM conducted the project administration. EM and DW were responsible for supervision. CM wrote the original draft. CM, CW, AR, CF, BM, NL, DW, and EM reviewed and edited the final manuscript.

All correspondence and requests for materials should be addressed to Esther Melamed.

## Ethics declarations / Competing interests

CM, CW, AR, CF, BM, NL, and DW have nothing to disclose. EM has received research funding from Babson Diagnostics, honorarium from Multiple Sclerosis Association of America and has served on advisory boards of Genentech, Horizon, Teva and Viela Bio.

## References

1. Tortorella, D., Gewurz, B. E., Furman, M. H., Schust, D. J. & Ploegh, H. L. Viral subversion of the immune system. Annu Rev Immunol 18, 861–926 (2000).

2. Gowthaman, U. & Eswarakumar, V. P. Molecular mimicry. Virulence 4, 433–434 (2013).

3. Chen, J. W. et al. Positive and negative selection shape the human naive B cell repertoire. J Clin Invest 132, e150985 (2022).

4. Palmer, E. Negative selection--clearing out the bad apples from the T-cell repertoire. Nat Rev Immunol 3, 383–391 (2003).

5. Cusick, M. F., Libbey, J. E. & Fujinami, R. S. Molecular mimicry as a mechanism of autoimmune disease. Clin Rev Allergy Immunol 42, 102–111 (2012).

6. Smatti, M. K. et al. Viruses and Autoimmunity: A Review on the Potential Interaction and Molecular Mechanisms. Viruses 11, 762 (2019).

7. Zhao, Z.-S., Granucci, F., Yeh, L., Schaffer, P. A. & Cantor, H. Molecular Mimicry by Herpes Simplex Virus-Type 1: Autoimmune Disease After Viral Infection. Science 279, 1344–1347 (1998).

8. Sabbatini, A., Bombardieri, S. & Migliorini, P. Autoantibodies from patients with systemic lupus erythematosus bind a shared sequence of SmD and Epstein-Barr virus-encoded nuclear antigen EBNA I. Eur J Immunol 23, 1146–1152 (1993).

9. Lasso, G., Honig, B. & Shapira, S. D. A Sweep of Earth’s Virome Reveals Host-Guided Viral Protein Structural Mimicry and Points to Determinants of Human Disease. Cell Systems 12, 82–91.e3 (2021).

10. Chang, S. T., Ghosh, D., Kirschner, D. E. & Linderman, J. J. Peptide length-based prediction of peptide–MHC class II binding. Bioinformatics 22, 2761–2767 (2006).

11. Wieczorek, M. et al. Major Histocompatibility Complex (MHC) Class I and MHC Class II Proteins: Conformational Plasticity in Antigen Presentation. Front Immunol 8, 292 (2017).

12. Trolle, T. et al. The Length Distribution of Class I-Restricted T Cell Epitopes Is Determined by Both Peptide Supply and MHC Allele-Specific Binding Preference. J Immunol 196, 1480– 1487 (2016).

13. Burrows, S. R., Rossjohn, J. & McCluskey, J. Have we cut ourselves too short in mapping CTL epitopes? Trends in Immunology 27, 11–16 (2006).

14. Buus, S. et al. High-resolution mapping of linear antibody epitopes using ultrahigh-density peptide microarrays. Mol Cell Proteomics 11, 1790–1800 (2012).

15. Qi, H. et al. Antibody Binding Epitope Mapping (AbMap) of Hundred Antibodies in a Single Run. Mol Cell Proteomics 20, 100059 (2021).

16. Doxey, A. C. & McConkey, B. J. Prediction of molecular mimicry candidates in human pathogenic bacteria. Virulence 4, 453–466 (2013).

17. Lebeau, G. et al. Zika E Glycan Loop Region and Guillain–Barré Syndrome-Related Proteins: A Possible Molecular Mimicry to Be Taken in Account for Vaccine Development. Vaccines 9, 283 (2021).

18. Adiguzel, Y. Molecular mimicry between SARS-CoV-2 and human proteins. Autoimmun Rev 20, 102791 (2021).

19. Begum, S. et al. Molecular Mimicry Analyses Unveiled the Human Herpes Simplex and Poxvirus Epitopes as Possible Candidates to Incite Autoimmunity. Pathogens 11, 1362 (2022).

20. Hurford, A. & Day, T. Immune evasion and the evolution of molecular mimicry in parasites. Evolution 67, 2889–2904 (2013).

21. Masson, P. et al. ViralZone: recent updates to the virus knowledge resource. Nucleic Acids Res 41, D579–583 (2013).

22. Schoch, C. L. et al. NCBI Taxonomy: a comprehensive update on curation, resources and tools. Database (Oxford) 2020, baaa062 (2020).

23. Kanehisa, M. & Goto, S. KEGG: kyoto encyclopedia of genes and genomes. Nucleic Acids Res 28, 27–30 (2000).

24. Uhlén, M. et al. Proteomics. Tissue-based map of the human proteome. Science 347, 1260419 (2015).

25. Gabrielsen, I. S. M. et al. Transcriptomes of antigen presenting cells in human thymus. PLoS One 14, e0218858 (2019).

26. Zamecnik, C. R. et al. A Predictive Autoantibody Signature in Multiple Sclerosis. medRxiv 2023.05.01.23288943 (2023) doi:10.1101/2023.05.01.23288943.

27. Bezanson, J., Edelman, A., Karpinski, S. & Shah, V. B. Julia: A fresh approach to numerical computing. SIAM review 59, 65–98 (2017).

28. R Core Team. R: *A Language and Environment for Statistical Computing*. (R Foundation for Statistical Computing, Vienna, Austria, 2021).

29. UniProt Consortium. UniProt: the Universal Protein Knowledgebase in 2023. Nucleic Acids Res 51, D523–D531 (2023).

30. Kang, M.-S. & Kieff, E. Epstein–Barr virus latent genes. Exp Mol Med 47, e131 (2015).

31. Slobedman, B., Barry, P. A., Spencer, J. V., Avdic, S. & Abendroth, A. Virus-encoded homologs of cellular interleukin-10 and their control of host immune function. J Virol 83, 9618–9629 (2009).

32. Xing, Y. & Hogquist, K. A. T-cell tolerance: central and peripheral. Cold Spring Harb Perspect Biol 4, a006957 (2012).

33. Bjornevik, K. et al. Longitudinal analysis reveals high prevalence of Epstein-Barr virus associated with multiple sclerosis. Science 375, 296–301 (2022).

34. Hrycek, A., Kuśmierz, D., Mazurek, U. & Wilczok, T. Human cytomegalovirus in patients with systemic lupus erythematosus. Autoimmunity 38, 487–491 (2005).

35. Israeli, E., Agmon-Levin, N., Blank, M., Chapman, J. & Shoenfeld, Y. Guillain-Barré syndrome--a classical autoimmune disease triggered by infection or vaccination. Clin Rev Allergy Immunol 42, 121–130 (2012).

36. Oikarinen, M. et al. Enterovirus Infections Are Associated With the Development of Celiac Disease in a Birth Cohort Study. Front Immunol 11, 604529 (2020).

37. Yazdanpanah, N. & Rezaei, N. Autoimmune complications of COVID-19. J Med Virol 94, 54–62 (2022).

38. Gómez-Rial, J., Rivero-Calle, I., Salas, A. & Martinón-Torres, F. Rotavirus and autoimmunity. J Infect 81, 183–189 (2020).

39. Banko, A., Miljanovic, D. & Cirkovic, A. Systematic review with meta-analysis of active herpesvirus infections in patients with COVID-19: Old players on the new field. Int J Infect Dis 130, 108–125 (2023).

40. Baxby, D. Poxviruses. in Medical Microbiology (ed. Baron, S.) (University of Texas Medical Branch at Galveston, Galveston (TX), 1996).

41. Efridi, W. & Lappin, S. L. Poxviruses. in StatPearls (StatPearls Publishing, Treasure Island (FL), 2024).

42. Amrani, A. et al. Expansion of the antigenic repertoire of a single T cell receptor upon T cell activation. J Immunol 167, 655–666 (2001).

43. Zamvil, S. S., Spencer, C. M., Baranzini, S. E. & Cree, B. A. C. The Gut Microbiome in Neuromyelitis Optica. Neurotherapeutics 15, 92–101 (2018).

44. Lanz, T. V. et al. Clonally expanded B cells in multiple sclerosis bind EBV EBNA1 and GlialCAM. Nature 603, 321–327 (2022).

45. Rojas, J. M., Avia, M., Martín, V. & Sevilla, N. IL-10: A Multifunctional Cytokine in Viral Infections. J Immunol Res 2017, 6104054 (2017).

46. Albarnaz, J. D. et al. Molecular mimicry of NF-κB by vaccinia virus protein enables selective inhibition of antiviral responses. Nat Microbiol 7, 154–168 (2022).

47. Nantes University Hospital. An Open Single-Center, Phase I Proof of Concept Trial to Assess the Safety and Feasibility of Adoptive Cell Therapy With Autologous EBV-Specific Cytotoxic T Lymphocytes (CTL) in Patients With a First Clinical Episode Highly Suggestive of Multiple Sclerosis. https://clinicaltrials.gov/study/NCT02912897 (2023).

48. Michael, L. Effects of Antiviral Therapies on Epstein-Barr Virus Replication. https://clinicaltrials.gov/study/NCT05957913 (2023).

