## Supplemental Figures for "Molecular Mimicry as a Mechanism of Viral Immune Evasion and Autoimmunity"

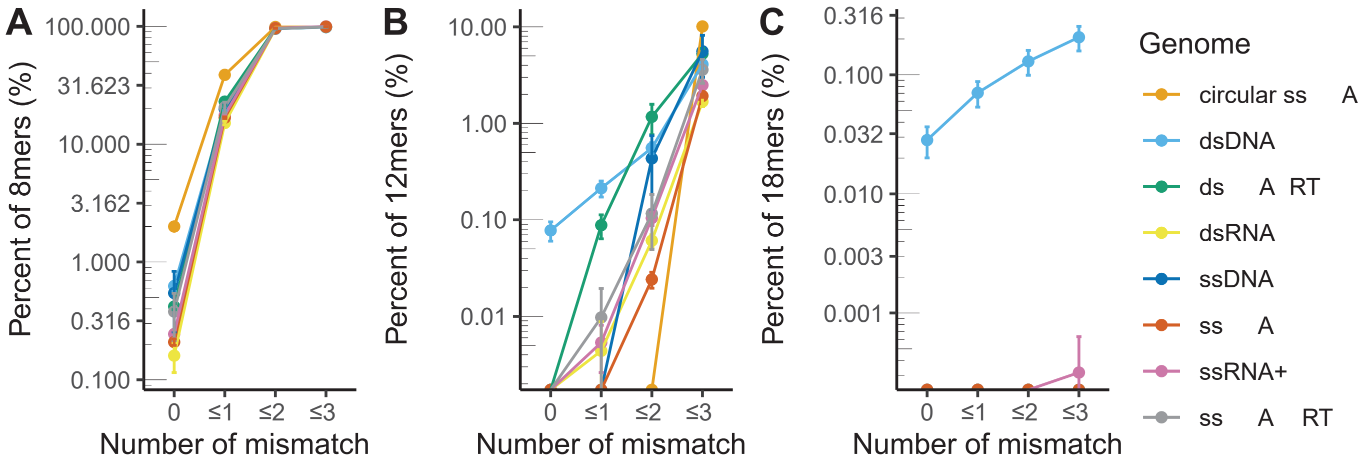
A
B
C
Percent of 8mers (%)
100.000
31.623
10.000
3.162
1.000
0.316
0.100
Percent of 12mers (%)
10.00
1.00
0.10
0.01
Percent of 18mers (%)
0.316
0.100
0.032
0.010
0.003
0.001
0
≤1
≤2
≤3
Number of mismatch
Genome
circular ss A
dsDNA
ds A RT
dsRNA
ssDNA
ss A
ssRNA+
ss A RT

**Figure S1 (Associated with Figure 1) – Evaluation of mimicry candidates by Baltimore classification.** Percent of **A)** 8mer, **B)** 12mers, and **C)** 18mers with 0, ≤1, ≤2, or ≤3 mismatches by Baltimore genome classification.

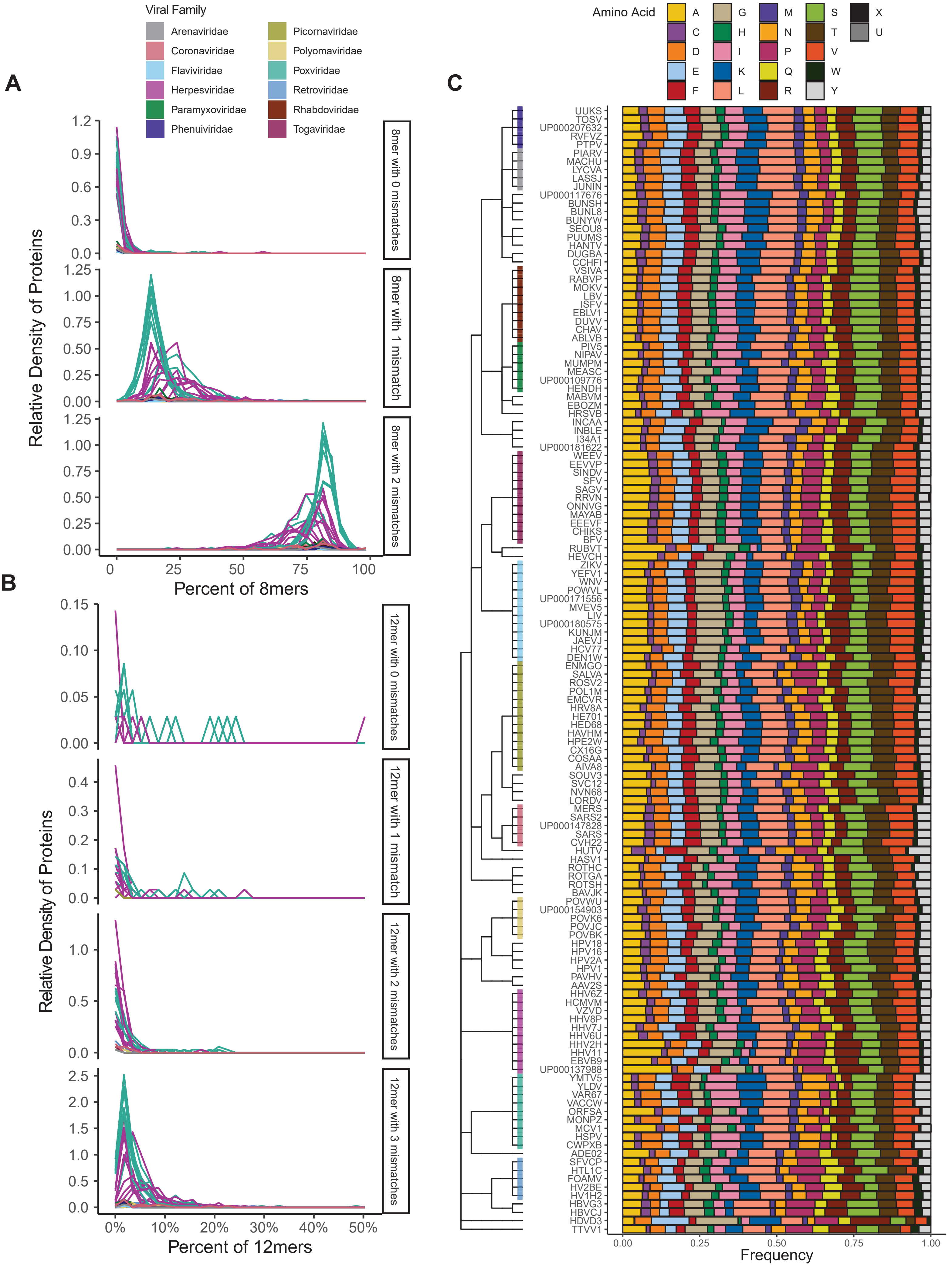

A
B
C
Viral Family
Arenaviridae
Coronaviridae
Flaviviridae
Herpesviridae
Paramyxoviridae
Phenuiviridae
Picornaviridae
Polyomaviridae
Poxviridae
Retroviridae
Rhabdoviridae
Togaviridae
Relative Density of Proteins
8mer with 0 mismatches
8mer with 1 mismatch
8mer with 2 mismatches
Percent of 8mers
12mer with 0 mismatches
12mer with 1 mismatch
12mer with 2 mismatches
12mer with 3 mismatches
Percent of 12mers
Amino Acid
A C D E F G H I K L M N P Q R S T V W Y X U
Frequency
UUKS
TOSV
UP000207632
RVFVZ
PTPV
PIARV
MACHU
LYCVA
LASSJ
JUNIN
UP000117676
BUNSH
BUNL8
BUNYW
SEOU8
PUUMS
HANTV
DUGBA
CCHFI
VSIVA
RABVP
MOKV
LBV
ISFV
EBLV1
DUVV
CHAV
ABLVB
PIV5
NIPAV
MUMPM
MEASC
UP000109776
HENDH
MABVM
EBOZM
HRSVB
INCAA
INBLE
I34A1
UP000181622
WEEV
EEVVP
SINDV
SFV
SAGV
RRVN
ONNVG
MAYAB
EEEVF
CHIKS
BFV
RUBVT
HEVCH
ZIKV
YEFV1
WNV
POWVL
UP000171556
MVEV5
LIV
UP000180575
KUNJM
JAEVJ
HCV77
DEN1W
ENMGO
SALVA
ROSV2
POL1M
EMCVR
HRV8A
HE701
HED68
HAVHM
HPE2W
CX16G
COSAA
AIVA8
SOUV3
SVC12
NVN68
LORDV
MERS
SARS2
UP000147828
SARS
CVH22
HUTV
HASV1
ROTHC
ROTGA
ROTSH
BAVJK
POVWU
UP000154903
POVK6
POVJC
POVBK
HPV18
HPV16
HPV2A
HPV1
PAVHV
AAV2S
HHV6Z
HCMVM
VZVD
HHV8P
HHV7J
HHV6U
HHV2H
HHV11
EBVB9
UP000137988
YMTV5
YLDV
VAR67
VACCW
ORFSA
MONPZ
MCV1
HSPV
CWPXB
ADE02
SFVCP
HTL1C
FOAMV
HV2BE
HV1H2
HBVG3
HBVCJ
HDVD3
TTVV1

**Figure S2 (Associated with Figure 1) – Molecular mimicry is dispersed through the viral proteome.** For each virus (a line), the density of proteins that have varying percentages of either **A)** 8mers with 0, 1, or 2 mismatches or **B)** 12mers with 0, 1, 2, or 3 mismatches is plotted. Color indicates the family of the virus. As shown by the frequency plot peaking to the left, viruses overall display a pattern of low mimicry rates in any given protein. **C)** The relative proportion of amino acid usage for each virus organized by viral phylogeny. “U” indicates selenocysteine and “X” indicates an unknown amino acid.

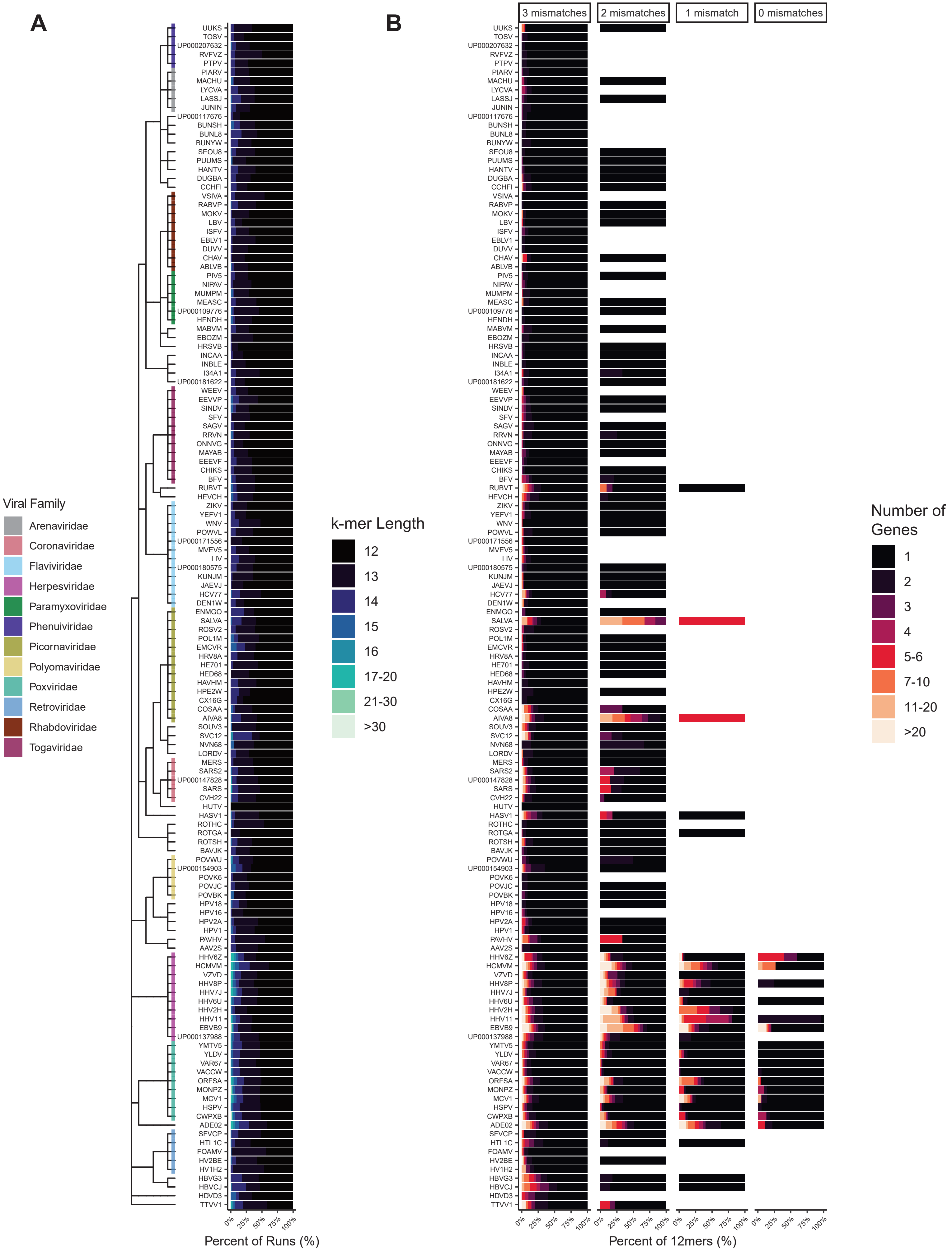
A
B
Viral Family
Arenaviridae
Coronaviridae
Flaviviridae
Herpesviridae
Paramyxoviridae
Phenuiviridae
Picornaviridae
Polyomaviridae
Poxviridae
Retroviridae
Rhabdoviridae
Togaviridae
k-mer Length
12
13
14
15
16
17-20
21-30
>30
Number of Genes
1
2
3
4
5-6
7-10
11-20
>20
3 mismatches
2 mismatches
1 mismatch
0 mismatches
UUKS
TOSV
UP000207632
RVFVZ
PTPV
PIARV
MACHU
LYCVA
LASSJ
JUNIN
UP000117676
BUNSH
BUNL8
BUNYW
SEOU8
PUUMS
HANTV
DUGBA
CCHFI
VSIVA
RABVP
MOKV
LBV
ISFV
EBLV1
DUVV
CHAV
ABLVB
PIV5
NIPAV
MUMPM
MEASC
UP000109776
HENDH
MABVM
EBOZM
HRSVB
INCAA
INBLE
I34A1
UP000181622
WEEV
EEVVP
SINDV
SFV
SAGV
RRVN
ONNVG
MAYAB
EEEVF
CHIKS
BFV
RUBVT
HEVCH
ZIKV
YEFV1
WNV
POWVL
UP000171556
MVEV5
LIV
UP000180575
KUNJM
JAEVJ
HCV77
DEN1W
ENMGO
SALVA
ROSV2
POL1M
EMCVR
HRV8A
HE701
HED68
HAVHM
HPE2W
CX16G
COSAA
AIVA8
SOUV3
SVC12
NVN68
LORDV
MERS
SARS2
UP000147828
SARS
CVH22
HUTV
HASV1
ROTHC
ROTGA
ROTSH
BAVJK
POVWU
UP000154903
POVK6
POVJC
POVBK
HPV18
HPV16
HPV2A
HPV1
PAVHV
AAV2S
HHV6Z
HCMVM
VZVD
HHV8P
HHV7J
HHV6U
HHV2H
HHV11
EBVB9
UP000137988
YMTV5
YLDV
VAR67
VACCW
ORFSA
MONPZ
MCV1
HSPV
CWPXB
ADE02
SFVCP
HTL1C
FOAMV
HV2BE
HV1H2
HBVG3
HBVCJ
HDVD3
TTVV1
0%
25%
50%
75%
100%
Percent of Runs (%)
Percent of 12mers (%)

**Figure S3 (Associated with Figure 3) – Phylogenetic analysis of run length and multi-mapping. A)** Percent of k-mers at various lengths for all viruses in the cohort organized by viral phylogeny. **B)** Percent of viral 12mers and their corresponding number of mimicked human genes under mismatch conditions of 0, ≤1, ≤2, and ≤3 mismatches.

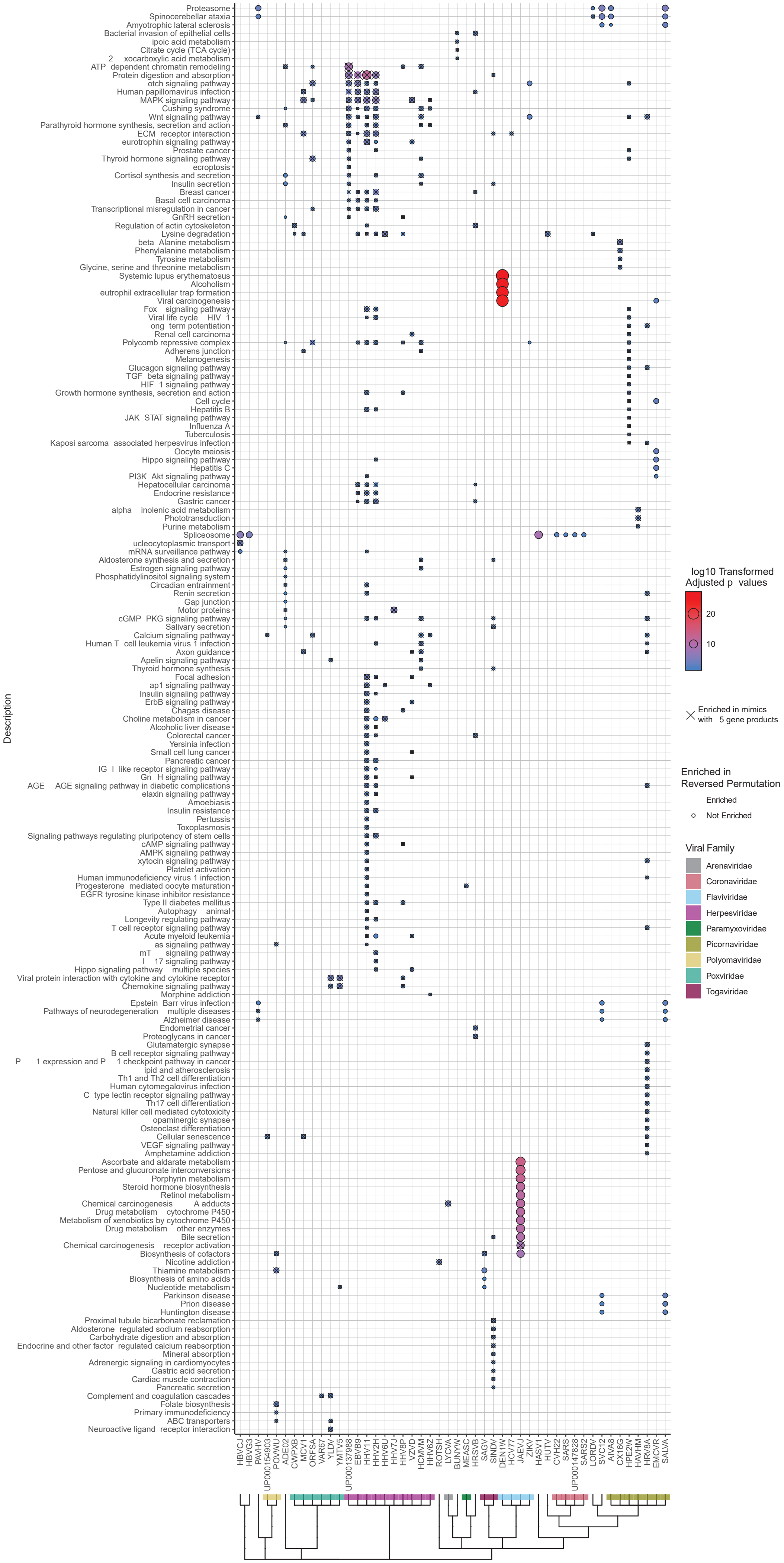
Proteasome
Spinocerebellar ataxia
Amyotrophic lateral sclerosis
Bacterial invasion of epithelial cells
ipoic acid metabolism
Citrate cycle (TCA cycle)
2 xocarboxylic acid metabolism
ATP dependent chromatin remodeling
Protein digestion and absorption
otch signaling pathway
Human papillomavirus infection
MAPK signaling pathway
Cushing syndrome
Wnt signaling pathway
Parathyroid hormone synthesis, secretion and action
ECM receptor interaction
eurotrophin signaling pathway
Prostate cancer
Thyroid hormone signaling pathway
ecroptosis
Cortisol synthesis and secretion
Insulin secretion
Breast cancer
Basal cell carcinoma
Transcriptional misregulation in cancer
GnRH secretion
Regulation of actin cytoskeleton
Lysine degradation
beta Alanine metabolism
Phenylalanine metabolism
Tyrosine metabolism
Glycine, serine and threonine metabolism
Systemic lupus erythematosus
Alcoholism
eutrophil extracellular trap formation
Viral carcinogenesis
Fox signaling pathway
Viral life cycle HIV 1
ong term potentiation
Renal cell carcinoma
Polycomb repressive complex
Adherens junction
Melanogenesis
Glucagon signaling pathway
TGF beta signaling pathway
HIF 1 signaling pathway
Growth hormone synthesis, secretion and action
Cell cycle
Hepatitis B
JAK STAT signaling pathway
Influenza A
Tuberculosis
Kaposi sarcoma associated herpesvirus infection
Oocyte meiosis
Hippo signaling pathway
Hepatitis C
PI3K Akt signaling pathway
Hepatocellular carcinoma
Endocrine resistance
Gastric cancer
alpha inolenic acid metabolism
Phototransduction
Purine metabolism
Spliceosome
ucleocytoplasmic transport
mRNA surveillance pathway
Aldosterone synthesis and secretion
Estrogen signaling pathway
Phosphatidylinositol signaling system
Circadian entrainment
Renin secretion
Gap junction
Motor proteins
cGMP PKG signaling pathway
Salivary secretion
Calcium signaling pathway
Human T cell leukemia virus 1 infection
Axon guidance
Apelin signaling pathway
Thyroid hormone synthesis
Focal adhesion
ap1 signaling pathway
Insulin signaling pathway
ErbB signaling pathway
Chagas disease
Choline metabolism in cancer
Alcoholic liver disease
Colorectal cancer
Yersinia infection
Small cell lung cancer
Pancreatic cancer
IG I like receptor signaling pathway
Gn H signaling pathway
AGE AGE signaling pathway in diabetic complications
elaxin signaling pathway
Amoebiasis
Insulin resistance
Pertussis
Toxoplasmosis
Signaling pathways regulating pluripotency of stem cells
cAMP signaling pathway
AMPK signaling pathway
xytocin signaling pathway
Platelet activation
Human immunodeficiency virus 1 infection
Progesterone mediated oocyte maturation
EGFR tyrosine kinase inhibitor resistance
Type II diabetes mellitus
Autophagy animal
Longevity regulating pathway
T cell receptor signaling pathway
Acute myeloid leukemia
as signaling pathway
mT signaling pathway
I 17 signaling pathway
Hippo signaling pathway multiple species
Viral protein interaction with cytokine and cytokine receptor
Chemokine signaling pathway
Morphine addiction
Epstein Barr virus infection
Pathways of neurodegeneration multiple diseases
Alzheimer disease
Endometrial cancer
Proteoglycans in cancer
Glutamatergic synapse
B cell receptor signaling pathway
P 1 expression and P 1 checkpoint pathway in cancer
ipid and atherosclerosis
Th1 and Th2 cell differentiation
Human cytomegalovirus infection
C type lectin receptor signaling pathway
Th17 cell differentiation
Natural killer cell mediated cytotoxicity
opaminergic synapse
Osteoclast differentiation
Cellular senescence
VEGF signaling pathway
Amphetamine addiction
Ascorbate and aldarate metabolism
Pentose and glucuronate interconversions
Porphyrin metabolism
Steroid hormone biosynthesis
Retinol metabolism
Chemical carcinogenesis A adducts
Drug metabolism cytochrome P450
Metabolism of xenobiotics by cytochrome P450
Drug metabolism other enzymes
Bile secretion
Chemical carcinogenesis receptor activation
Biosynthesis of cofactors
Nicotine addiction
Thiamine metabolism
Biosynthesis of amino acids
Nucleotide metabolism
Parkinson disease
Prion disease
Huntington disease
Proximal tubule bicarbonate reclamation
Aldosterone regulated sodium reabsorption
Carbohydrate digestion and absorption
Endocrine and other factor regulated calcium reabsorption
Mineral absorption
Adrenergic signaling in cardiomyocytes
Gastric acid secretion
Cardiac muscle contraction
Pancreatic secretion
Complement and coagulation cascades
Folate biosynthesis
Primary immunodeficiency
ABC transporters
Neuroactive ligand receptor interaction
Description
HBVCJ
HBVG3
PAVHV
UP000154903
POVWU
ADE02
CWPXB
MCV1
ORFSA
VAR67
YLDV
YMTV5
UP000137988
EBVB9
HHV11
HHV2H
HHV6U
HHV7J
HHV8P
VZVD
HCMVM
HHV6Z
ROTSH
LYCVA
BUNYW
MEASC
HRSVB
SAGV
SINDV
DEN1W
HCV77
JAEVJ
ZIKV
HASV1
HUTV
CVH22
SARS
UP000147828
SARS2
LORDV
SVC12
AIVA8
CX16G
HPE2W
HAVHM
HRV8A
EMCVR
SALVA
log10 Transformed
Adjusted p values
20
10
Enriched in mimics
with 5 gene products
Enriched in
Reversed Permutation
Enriched
Not Enriched
Viral Family
Arenaviridae
Coronaviridae
Flaviviridae
Herpesviridae
Paramyxoviridae
Picornaviridae
Polyomaviridae
Poxviridae
Togaviridae

**Figure S4 (Associated with Figure 4) – Complete enrichment results for all viruses.** Hypergeometric enrichment testing of KEGG pathways for human proteins that are mimicked by each virus. Significant enrichment is displayed as a dot and is outlined if that biological enrichment was not observed in the reverse proteome permutation (permutation 2). Viruses are organized by their phylogeny.

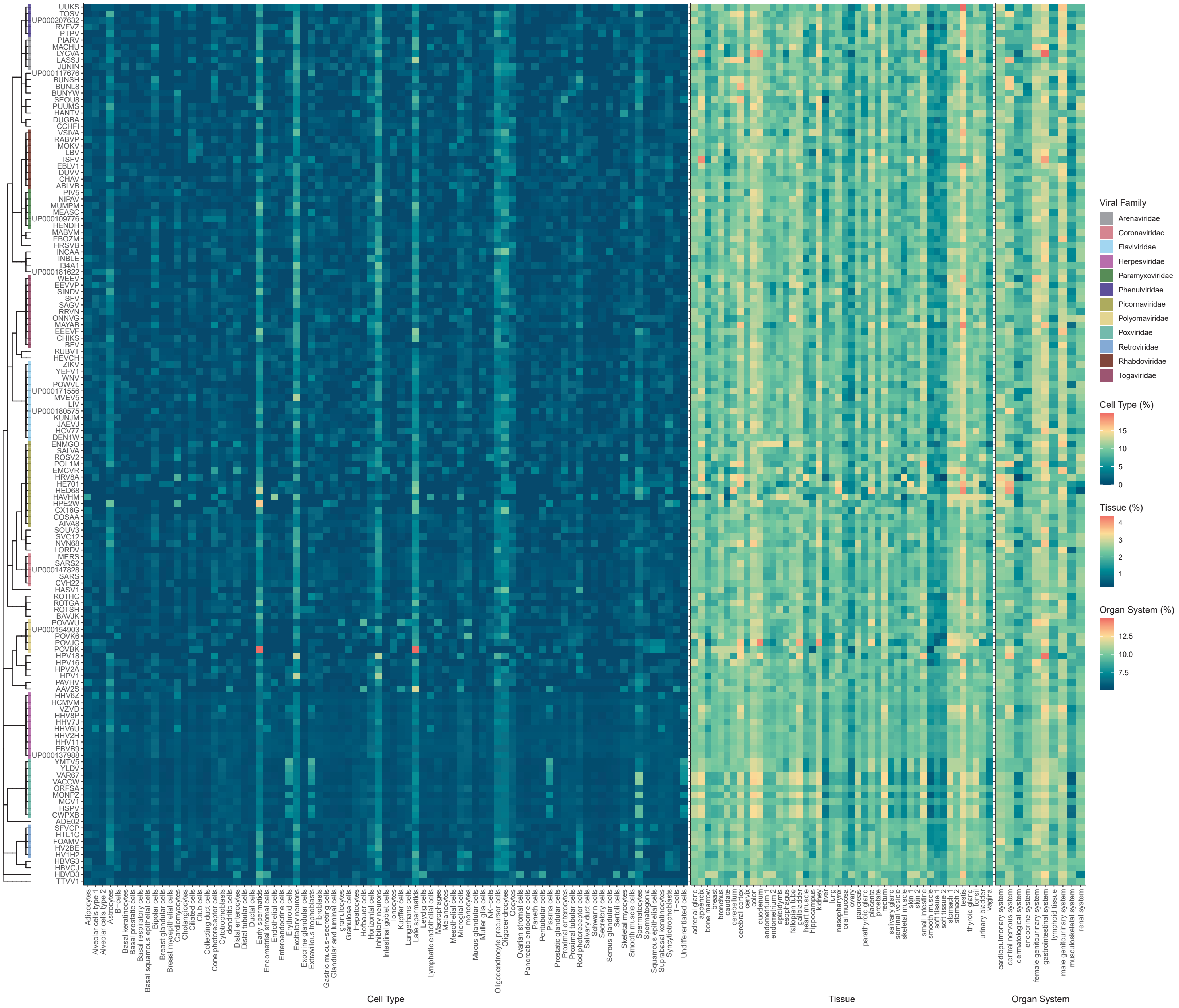

UUKS
TOSV
UP000207632
RVFVZ
PTPV
PIARV
MACHU
LYCVA
LASSJ
JUNIN
UP000117676
BUNSH
BUNL8
BUNYW
SEOU8
PUUMS
HANTV
DUGBA
CCHFI
VSIVA
RABVP
MOKV
LBV
ISFV
EBLV1
DUVV
CHAV
ABLVB
PIV5
NIPAV
MUMPM
MEASC
UP000109776
HENDH
MABVM
EBOZM
HRSVB
INCAA
INBLE
I34A1
UP000181622
WEEV
EEVVP
SINDV
SFV
SAGV
RRVN
ONNVG
MAYAB
EEEVF
CHIKS
BFV
RUBVT
HEVCH
ZIKV
YEFV1
WNV
POWVL
UP000171556
MVEV5
LIV
UP000180575
KUNJM
JAEVJ
HCV77
DEN1W
ENMGO
SALVA
ROSV2
POL1M
EMCVR
HRV8A
HE701
HED68
HAVHM
HPE2W
CX16G
COSAA
AIVA8
SOUV3
SVC12
NVN68
LORDV
MERS
SARS2
UP000147828
SARS
CVH22
HASV1
ROTHC
ROTGA
ROTSH
BAVJK
POVWU
UP000154903
POVK6
POVJC
POVBK
HPV18
HPV16
HPV2A
HPV1
PAVHV
AAV2S
HHV6Z
HCMVM
VZVD
HHV8P
HHV7J
HHV6U
HHV2H
HHV11
EBVB9
UP000137988
YMTV5
YLDV
VAR67
VACCW
ORFSA
MONPZ
MCV1
HSPV
CWPXB
ADE02
SFVCP
HTL1C
FOAMV
HV1H2
HBVG3
HBVCJ
HDVD3
TTVV1
Adipocytes
Alveolar cells type 1
Alveolar cells type 2
Astrocytes
B−cells
Basal keratinocytes
Basal prostatic cells
Basal respiratory cells
Basal squamous epithelial cells
Bipolar cells
Breast glandular cells
Breast myoepithelial cells
Cardiomyocytes
Cholangiocytes
Ciliated cells
Club cells
Collecting duct cells
Cone photoreceptor cells
Cytotrophoblasts
dendritic cells
Distal enterocytes
Distal tubular cells
Ductal cells
Early spermatids
Endometrial stromal cells
Endothelial cells
Enteroendocrine cells
Erythroid cells
Excitatory neurons
Exocrine glandular cells
Extravillous trophoblasts
Fibroblasts
Gastric mucus−secreting cells
Glandular and luminal cells
granulocytes
Granulosa cells
Hepatocytes
Hofbauer cells
Horizontal cells
Inhibitory neurons
Intestinal goblet cells
Ionocytes
Kupffer cells
Langerhans cells
Late spermatids
Leydig cells
Lymphatic endothelial cells
Macrophages
Melanocytes
Mesothelial cells
Microglial cells
monocytes
Mucus glandular cells
Muller glia cells
NK−cells
Oligodendrocyte precursor cells
Oligodendrocytes
Oocytes
Ovarian stromal cells
Pancreatic endocrine cells
Paneth cells
Peritubular cells
Plasma cells
Prostatic glandular cells
Proximal enterocytes
Proximal tubular cells
Rod photoreceptor cells
Salivary duct cells
Schwann cells
Secretory cells
Serous glandular cells
Sertoli cells
Skeletal myocytes
Smooth muscle cells
Spermatocytes
Spermatogonia
Squamous epithelial cells
Suprabasal keratinocytes
Syncytiotrophoblasts
T−cells
Undifferentiated cells
Cell Type
adrenal gland
appendix
bone marrow
breast
bronchus
caudate
cerebellum
cerebral cortex
cervix
colon
duodenum
endometrium 1
endometrium 2
epididymis
esophagus
fallopian tube
gallbladder
heart muscle
hippocampus
kidney
liver
lung
nasopharynx
oral mucosa
ovary
pancreas
parathyroid gland
placenta
prostate
rectum
salivary gland
seminal vesicle
skeletal muscle
skin 1
skin 2
small intestine
smooth muscle
soft tissue 1
soft tissue 2
stomach 1
stomach 2
testis
thyroid gland
tonsil
urinary bladder
vagina
Tissue
cardiopulmonary system
central nervous system
dermatological system
endocrine system
female genitourinary system
gastrointestinal system
lymphoid tissue
male genitourinary system
musculoskeletal system
renal system
Organ System
Viral Family
Arenaviridae
Coronaviridae
Flaviviridae
Herpesviridae
Paramyxoviridae
Phenuiviridae
Picornaviridae
Polyomaviridae
Poxviridae
Retroviridae
Rhabdoviridae
Togaviridae
Cell Type (%)
15
10
5
0
Tissue (%)
4
3
2
1
Organ System (%)
12.5
10.0
7.5

**Figure S5 (Associated with Figure 4) – Cell type, tissue, and organ system percentages of 12mers.** Heatmap of the percent of 12mer mimics with 3 or less mismatches that are known to be expressed in varying cell types, tissues, and organ systems as reported in the Human Protein Atlas. .

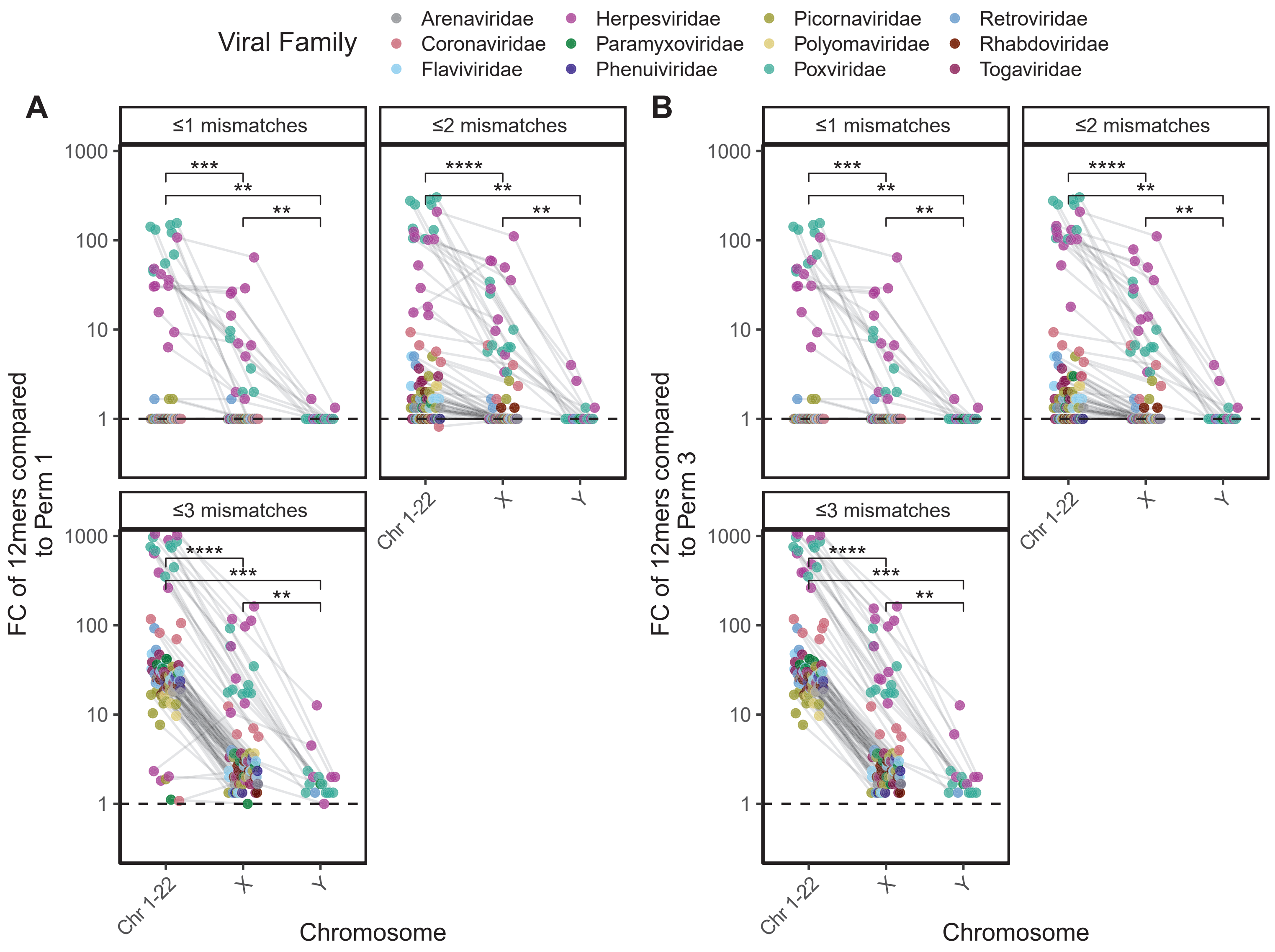
Viral Family
Arenaviridae
Coronaviridae
Flaviviridae
Herpesviridae
Paramyxoviridae
Phenuiviridae
Picornaviridae
Polyomaviridae
Poxviridae
Retroviridae
Rhabdoviridae
Togaviridae
A
B
≤1 mismatches
≤2 mismatches
≤3 mismatches
FC of 12mers compared to Perm 1
FC of 12mers compared to Perm 3
1000
100
10
1
Chr 1-22
X
Y
Chromosome

**Figure S6 (Associated with Figure 4) – Chromosome enrichment compared to A) permutation 1 and B) permutation 3.** Fold change of the percent of mimics whose human counterpart is encoded on either an autosome (Chromosomes 1-22), X, or Y chromosome over the rate in the **A)** randomly shuffled proteome (permutation 1) and **B)** the AA class shuffled proteome (permutation 3). Wilcoxon summed-rank test used for paired pairwise comparison. All p values adjusted for multiple hypothesis testing using Benjamini-Hochberg corrections (* p.adj ≤ 0.05, ** p.adj ≤ 0.01, *** p.adj ≤ 0.001, **** p.adj ≤ 0.0001).

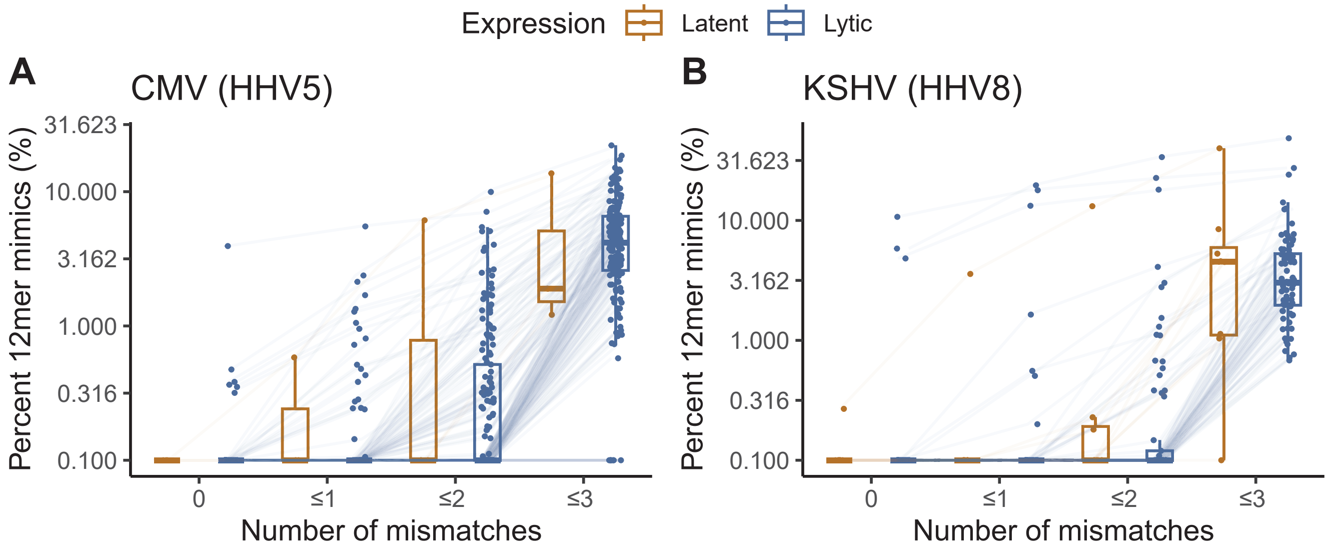

Expression
Latent
Lytic
A
CMV (HHV5)
B
KSHV (HHV8)
Percent 12mer mimics (%)
31.623
10.000
3.162
1.000
0.316
0.100
0
≤1
≤2
≤3
Number of mismatches

**Figure S7 (Associated with Figure 5) –** Percent of 12mers with 0, ≤1, ≤2, and ≤3 mismatches in latent and lytic **A)** CMV, and **B)** HHV8 proteins. Each point denotes a single protein connected across the mismatch levels with boxplots denoting the interquartile range. Wilcoxon summed-rank test used for paired pairwise comparison. All p values adjusted for multiple hypothesis testing using Benjamini-Hochberg corrections (* p.adj ≤ 0.05, ** p.adj ≤ 0.01, *** p.adj ≤ 0.001).

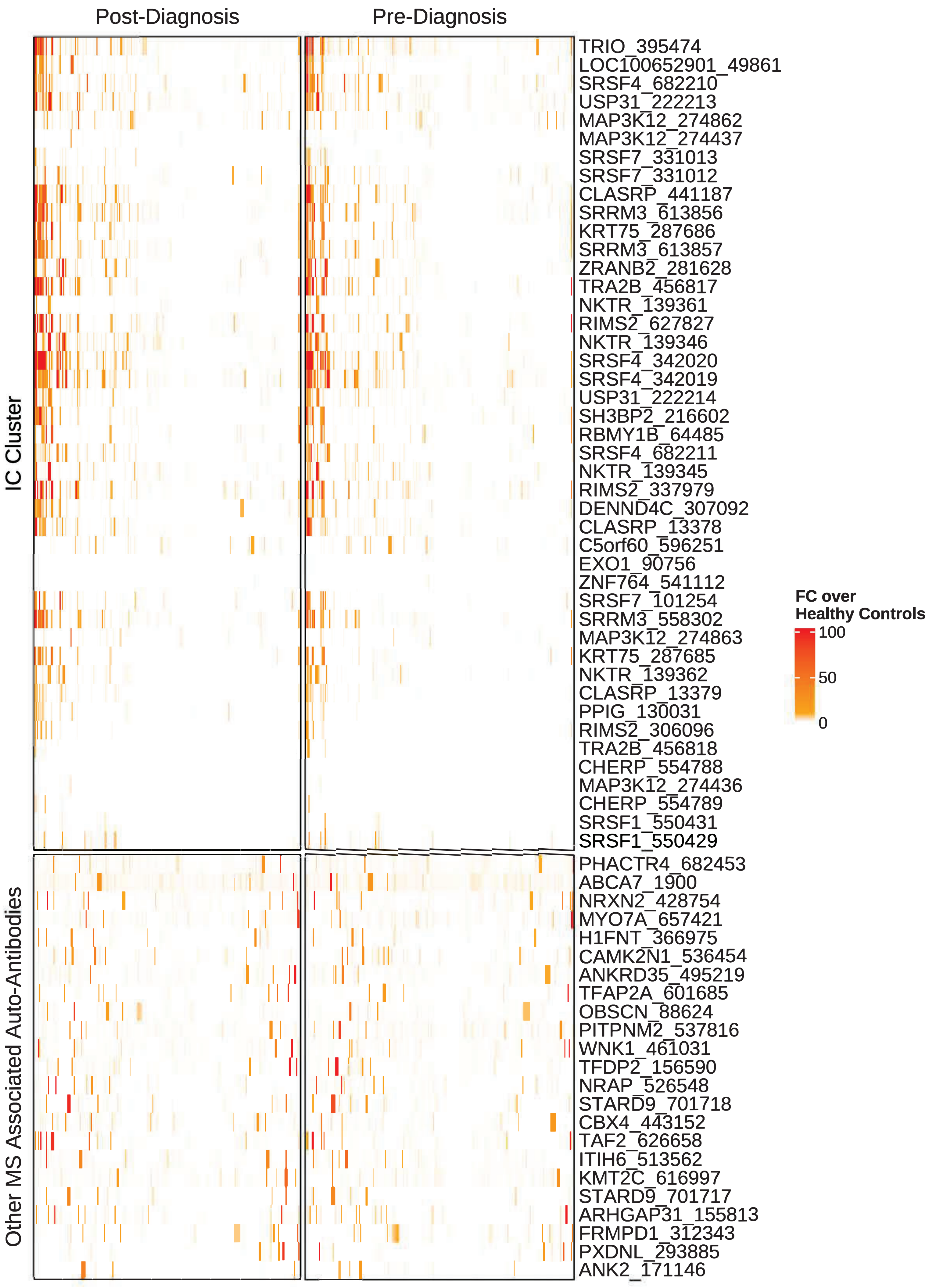
Post-Diagnosis
Pre-Diagnosis
IC Cluster
Other MS Associated Auto-Antibodies
TRIO_395474
LOC100652901_49861
SRSF4_682210
USP31_222213
MAP3K12_274862
MAP3K12_274437
SRSF7_331013
SRSF7_331012
CLASRP_441187
SRRM3_613856
KRT75_287686
SRRM3_613857
ZRANB2_281628
TRA2B_456817
NKTR_139361
RIMS2_627827
NKTR_139346
SRSF4_342020
SRSF4_342019
USP31_222214
SH3BP2_216602
RBMY1B_64485
SRSF4_682211
NKTR_139345
RIMS2_337979
DENND4C_307092
CLASRP_13378
C5orf60_596251
EXO1_90756
ZNF764_541112
SRSF7_101254
SRRM3_558302
MAP3K12_274863
KRT75_287685
NKTR_139362
CLASRP_13379
PPIG_130031
RIMS2_306096
TRA2B_456818
CHERP_554788
MAP3K12_274436
CHERP_554789
SRSF1_550431
SRSF1_550429
PHACTR4_682453
ABCA7_1900
NRXN2_428754
MYO7A_657421
H1FNT_366975
CAMK2N1_536454
ANKRD35_495219
TFAP2A_601685
OBSCN_88624
PITPNM2_537816
WNK1_461031
TFDP2_156590
NRAP_526548
STARD9_701718
CBX4_443152
TAF2_626658
ITIH6_513562
KMT2C_616997
STARD9_701717
ARHGAP31_155813
FRMPD1_312343
PXDNL_293885
ANK2_171146
FC over
Healthy Controls
100
50
0

**Figure S8 (Associated with Figure 6) – Additional MS associated auto-antibodies outside the IC cluster from Zamecnik et al.**[26]. Heatmap of fold change for auto-antibodies in MS patients before and after diagnosis over healthy control levels reveals that MS auto-antibodies outside the IC cluster identified in Zamecnik et al.[26] are not concordant.
